# JNK1 And Downstream Signalling Hubs Regulate Anxiety-Like Behaviours In A Zebrafish Larvae Phenotypic Screen

**DOI:** 10.1101/2024.01.09.574843

**Authors:** Ye Hong, Christel Sourander, Benjamin Hackl, Jedidiah S. Patton, Jismi John, Ilkka Paatero, Eleanor Coffey

**Affiliations:** Turku Bioscience, University of Turku and Åbo Akademi University, Tykistökatu 6, Turku 20520, Finland

## Abstract

Current treatments for anxiety and depression show limited efficacy in many patients indicating that research into new underlying mechanisms is needed. Inhibition of JNK1 has been shown to evoke an anxiolytic-and antidepressant-like phenotype in mice however the downstream effectors that elicit these behavioural effects are unknown. Here we employ a zebrafish (*D. Rerio*) larvae behavioural assay to identify an antidepressant-/anxiolytic-like phenotype based on 2759 measured stereotypic responses to clinically proven antidepressant and anxiolytic (AA) drugs. Employing machine learning, we classify an AA phenotype from behavioural features measured during and after a startle battery in fish exposed to AA drugs (fluoxetine, imipramine, diazepam, lithium chloride, ketamine). We demonstrate that structurally independent JNK inhibitors replicate the AA classification with high accuracy, consistent with findings in mice. We go on to identify signalling hubs downstream from JNK1 by comparing phosphoproteome data from wildtype and *Jnk1-/-* mouse brains, and test these hubs as possible mediators of the AA phenotype in zebrafish larvae. Among these, we find that AKT, GSK-3, 14-3-3ζ/ε and PKCε, when pharmacologically targeted, phenocopy clinically proven AA drugs. This assay shows promise as an early phase screening for compounds with anti-stress-axis/anxiolytic-like properties, and for mode of action analysis.

## INTRODUCTION

Anxiety and depression are highly prevalent mental disorders that involve abnormal neural function and dysregulation of circuits associated with threat and fear responses ^1,2^. Together they account for over half of the global burden of mental disorders, and are among the largest of disease burdens overall ^3,4^. Current treatments are lacking as not all patients achieve remission from their symptoms ^5^, while side effects are common ^6^. These disorders develop as a result of a complex interplay between genes and the environment ^7,8^, and involve a wide variety of mechanisms ^5,9^. One of the most consistent changes associated with depression is increased plasma cortisol and deregulation of hypothalamic-pituitary-axis homeostasis associated with desensitised cortisol receptor feedback mechanisms ^10,11^. Clinically useful antidepressant drugs such as serotonin reuptake inhibitors (SSRIs) and tricyclic antidepressants have been shown to recover cortisol homeostasis ^12^. A screening assay based on behaviour can potentially accelerate mode of action analysis and mechanistic understanding.

Phenotypic screening represents a strategy that can identify molecules with the ability to alter behaviour ^13^. While mouse models are used to model disease, the time and resources needed to screen compounds in mice would be prohibitive, thus simpler systems are needed when large numbers of compounds should be tested. The zebrafish larvae as a model organism is emerging as a viable approach for screening of phenotypic behaviours ^14–16^. For analysis of stress-induced behaviour, the zebrafish hypothalamic-pituitary-interrenal (HPI) axis is cortisol responsive already by four days post fertilization, and like the HP-adrenal (HPA) axis in mammals, its dysregulation is associated with dysfunctional coping behaviours ^17,18^. Also, zebrafish larva habenula and amygdala are involved in affective behaviours at an early stage ^17^, and zebrafish brain neurotransmitters and corresponding receptors (including NMDA receptors) are expressed and functional at the larval stage. Zebrafish express orthologues for about 70 % of human genes with 47 % of these genes bearing a 1:1 relationship ^19,20^, compared to 80 % in mice ^21^. The larvae display stereotypical behavioural responses such as thigmotaxis which is an evolutionarily conserved anxiety behavioural response ^22^, and hyper-locomotor activity in response to neuroactive drugs ^15,16,23^. For these reasons it is used for central nervous system drug screens at an early developmental stage which is suitable for scaleup ^18,24^.

The JNK1 pathway represents a potential regulator of anxiety and depression. In brain there are three *JNK* genes and 10 splice variants expressed. Among these, JNK1 is known to play important roles during morphogenesis and in axonal/dendritic architecture modelling during brain development ^25–28^. In nervous tissue, JNK has been shown to be activated by endocrine stress leading to retraction of dendritic spines, and JNK-inhibitor treated or *Jnk1-/-* mice show reduced anxiety- and depressive-like behaviours ^29–31^. Moreover, JNK signalling may contribute to glucocorticoid resistance and HPA-axis dysregulation underlying anxiety and depression ^32^, as JNK phosphorylation of the glucocorticoid receptor blocks its nuclear translocation and subsequent gene regulation ^33,34^. Also, JNK activity is reduced by the fast-acting anti-depressant ketamine when administered to hippocampal neurons in culture ^31^. However, JNK1 is expected to phosphorylate a large number of downstream targets in brain ^28^, and whether any of these downstream targets contribute to anxiety and depressive-like responses is not known.

In this study we identify signalling hubs downstream of JNK1 by comparing the mouse brain phosphoproteome from wild-type and *Jnk1-/-* mice. We then utilize a zebrafish larvae behavioural assay to test whether any of these downstream hubs phenocopy known antidepressant and anxiolytic drugs, and JNK1 inhibition. This identifies downstream players from JNK1 that phenocopy AA-like stereotypic responses in a zebrafish larvae screen.

## RESULTS

### Establishing an acoustic/light stimulus battery to profile zebrafish larvae behavioural responses to anti-depressant and anxiolytic drugs

We developed a multiparameter test battery to assess the behaviour of zebrafish larvae in response to light and acoustic stressors. For this, larvae were placed individually in square wells of a 96-well plate and exposed to blue (470 nm), red (635 nm), purple (a combination of red and blue lights), white light, or flickering light, combined with light or heavy tapping as indicated (Fig. 1A and B). Zebrafish larvae exhibited robust motility from 4 days post-fertilization (dpf) onwards for all stimuli, except for white light where older larvae were less responsive (Fig. 1C). We expanded the test to include five cycles, each followed by a 29-minute intermission post startle (Fig. 1D). This enabled us to monitor stress, recovery, and habituation responses.

**Figure 1.**
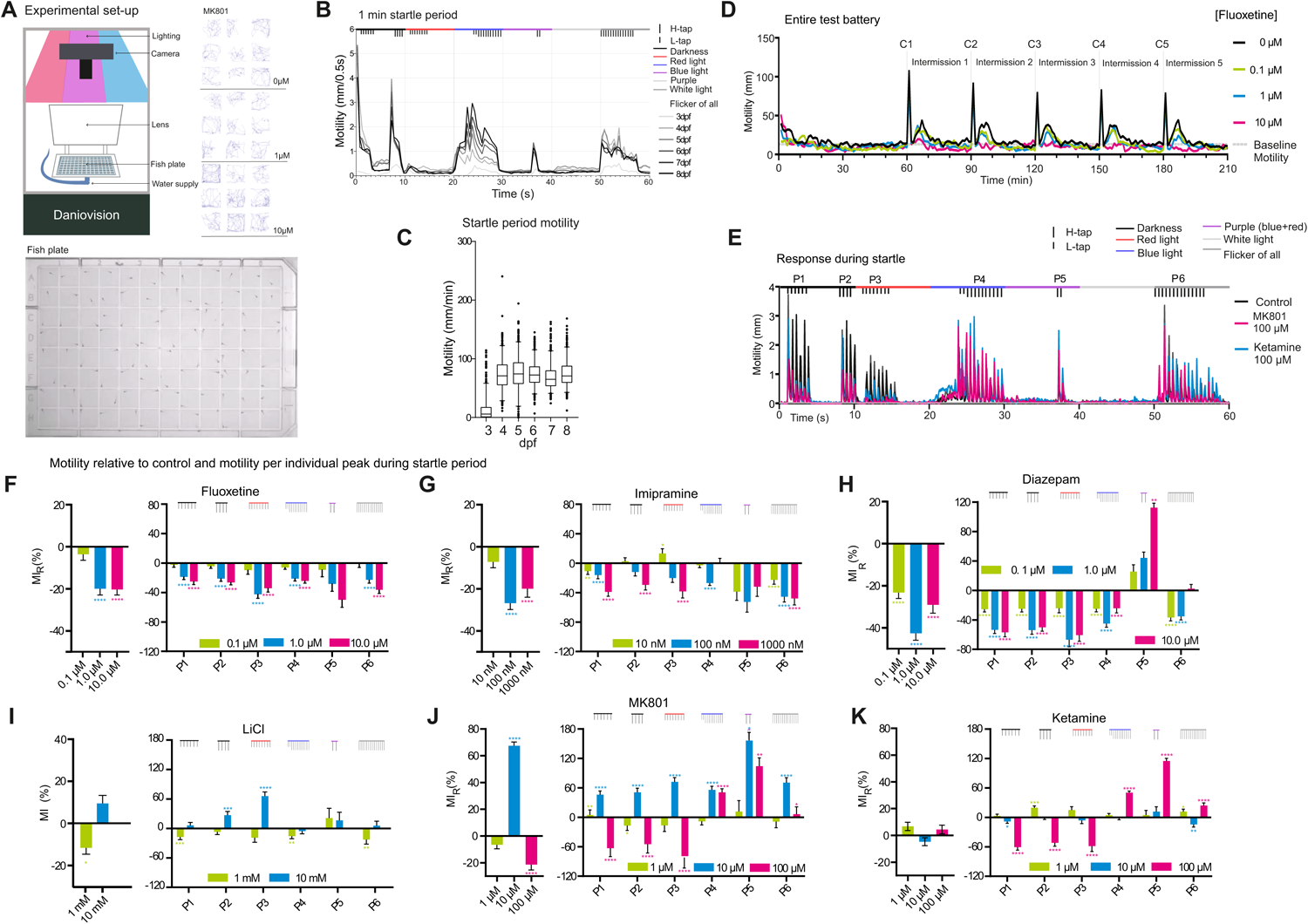
Motility profiles for anxiolytic and anti-depressant drugs. (A) A schematic of experimental setup shows the apparatus, square-well plate and representative motility traces from 1min tests from larvae treated with MK801. (B) Motility responses from zebrafish larvae at 3 to 8 days post fertilization (dpf) during the 1 min startle battery. (C) Mean motility data of zebrafish larvae at 3-8 dpf. Larvae number per age group; 3 dpf (n=95), 4 dpf (n=94), 5 dpf (n=94), 6 dpf (n=89), 7 dpf (n= 89) and 8 dpf (n=97). (D) Example traces from the entire test battery showing mean motility of 7 dpf fish. 60 min acclimation is followed by 5 cycles (C1 to C5) of 1 min startle period with 29 min recovery between cycles. Measurements are from larvae treated with or without fluoxetine (0.1, 1.0, or 10 µM) for 1 h (n=22 to 24). (E) A trace of 1 min startle response shows motility of fish treated with high doses of ketamine or MK801 (100 µM; n=115). (F-K) Motility responses are shown for AA drugs according to stimuli clusters P1 to P6. The left-side panels depict the overall mean motility during the 5 cycles of 1 min startle. The right-side panels show the mean motility data within the peaks only. Y-axis scaling differs to accommodate various size effects. One-way ANOVA with post-hoc Tukey test was performed on the original MI distributions to calculate the p-values of the left-side panels; Two-way ANOVA with post-hoc Dunnett test was performed on the original MI distributions to calculate the p-values of the right-side panels. * p-value≤0.05; ** p-value *≤ 0.01; ***p-value≤0.001; ****p-value≤0.0001. The number of fish measured for each treatment were as follows (the total number of repeats is shown in parenthesis): control: 136 (680 observations), diazepam: 70 (350 observations), fluoxetine: 72 (360 observations), imipramine: 69 (345 observations), LiCl: 41 (205 observations), ketamine: 70 (350 observations), MK801: 70 (350 observations).

### Classical and fast acting anti-depressant and anxiolytic drugs alter zebrafish larvae motility

We next measured motility of zebrafish larvae following a 1-hour incubation with classical <u>a</u>nti-depressant or <u>a</u>nxiolytic (AA) drugs: fluoxetine (a SSRI), imipramine (an antagonist against the serotonin transporter > noradrenaline transporter > dopamine transporter), LiCl (which targets inositol monophosphatase and GSK3 ^35^), diazepam (a Gamma Amino Butyric Acid receptor antagonist) or low dose (1 – 10 µM) ketamine; a fast-acting antidepressant, or high dose (100 µM) ketamine or MK801 (also known as dizocilpine)^36^; NMDA receptor antagonists with psychotic activities (Fig. 1E-K; Table 1). Additional data for each drug treatment (i.e. individual fish tracking plots and quantitative output from individual cycles) is found in supplementary figures 1 to 6. The responses to the anti-psychotic drug Haloperidol responses are shown for comparison with the AA drugs (Supplementary figure 7). Motility responses during the 1-minute startle period were quantified from six peak clusters (P1 to P6) (Fig. 1E).

**Table 1:**
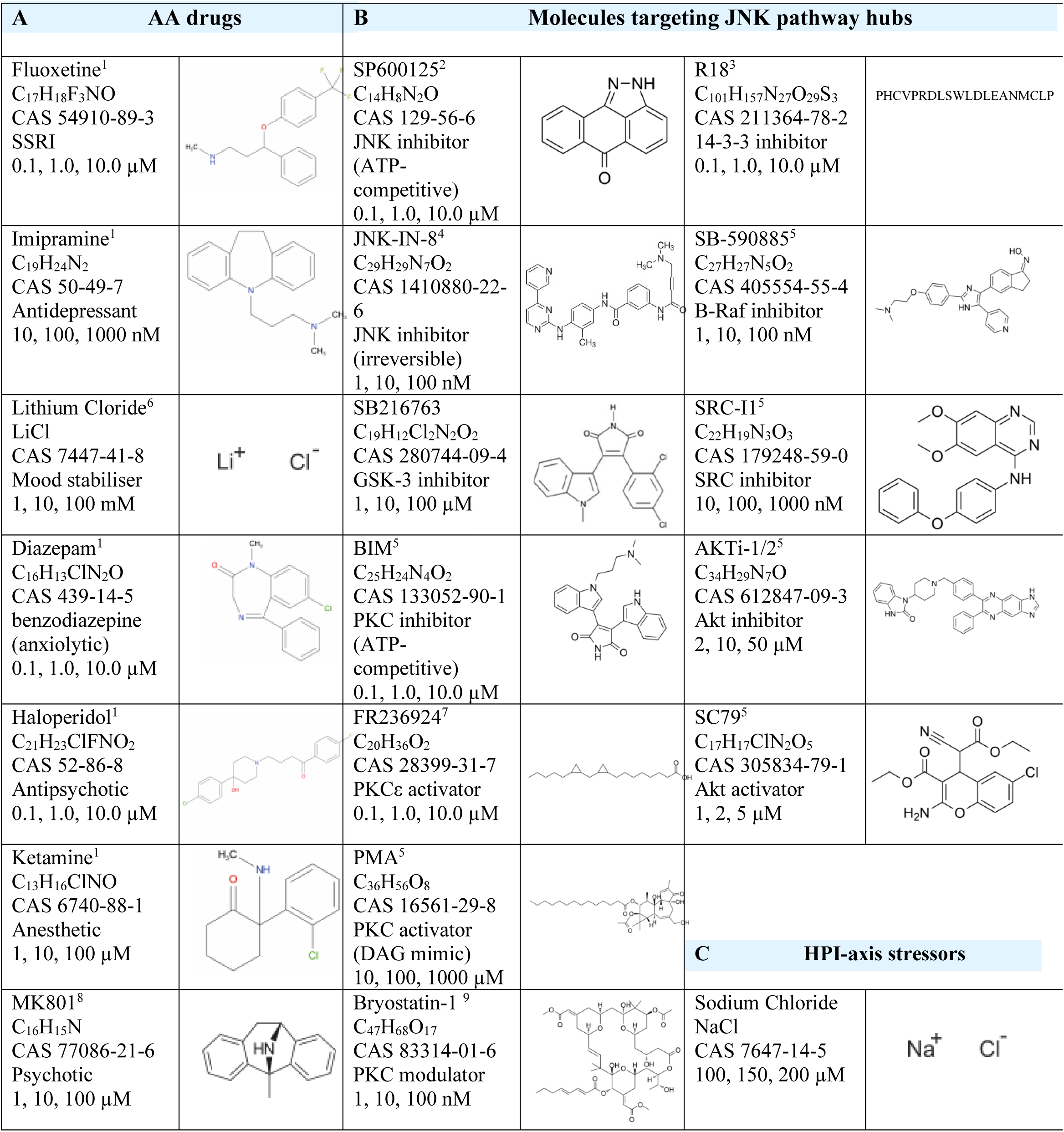
Drugs used in the study. Name, formula, CAS number, usage, used concentration and chemical structures are shown for (A) the antidepressant/anxiolytic (AA) drug group; (B) molecules targeting the JNK pathway hubs and (C) the known HPI axis stressors. (1) Wishart, D. S. *et al.* DrugBank 5.0 *Nucleic Acids Res* **46**, D1074-D1082 (2018). (2) Bennett, B. L. *et al. Proc Natl Acad Sci U S A* **98**, 13681-13686 (2001). (3) Wang, B. *et al.* I *Biochemistry* **38**, 12499-12504 (1999). (4) Zhang, T. *et al. Chem Biol* **19**, 140-154 (2012). (5) Ursu, O. *et al. Nucleic Acids Res* **45**, D932-D939 (2017). (6) Gould, T. D. & Manji, H. K. *Neuropsychopharmacology* **30**, 1223-1237 (2005). (7) Tanaka, A. & Nishizaki, T. *Bioorg Med Chem Lett* **13**, 1037-1040 (2003). (8) Huettner, J. E. & Bean, B. P. (9) Keenan, C., Goode, N. & Pears, C. *FEBS Lett* **415**, 101-108 (1997).

Clinically relevant doses of the AA drugs reduced the distance moved during the startle period (Fig. 1F-K). Even lithium at 1 mM showed a response similar to other AA drugs, while at 10 mM (above the therapeutic range), LiCl increased distance moved, possibly due to engagement of additional targets. Diazepam, with its muscle relaxant properties, caused a more significant motility inhibition than other AA drugs. However, it induced an unexpected increase in motility in response to purple light plus tapping (P5) (Fig. 1H), resembling the paradoxical hyperactivity or agitation effect seen in some individuals in response to diazepam ^16,37^. Overall the motility profiles from the AA drugs (at the clinically relevant dose) were similar.

Ketamine elicits an antidepressant effect in humans at steady state plasma levels of around 1 µM, and an anaesthetic effect at an average dose of 9.3 µM in humans ^38,39^. We therefore treated larvae with doses in this range. During the acute startle phase, ketamine decreased motility at doses of 10 µM and above for peaks P1-P3 and increased motility for peaks P4-P6. This closely matched the effect of MK801 and is therefore likely to be mediated through an NMDA receptor antagonist effect as both drugs share the phencyclidine site on (Fig. 1J-K). The response at lower doses of MK801 is consistent with the higher affinity of MK801 for NMDA receptor binding (Ki = 30.5 nM for MK801 verses 417 nM for ketamine) ^38^. In summary, treatment of zebrafish larvae with classical AA drugs suppressed their motility responses during the test battery, at doses that are clinically relevant for an anti-depressant/anxiolytic effect in mammals.

### Distance, turning, pausing and spurting are altered by AA drugs during the startle period

We next examined the behavioural syllables that make up the motility response. Thus, in addition to distance travelled, we also measured turning (sum of turning angles), pausing (total time spent pausing), spurt velocity, time thigmotaxis and distance thigmotaxis near the tank border (Fig. 2A). We characterised the effect of AA drugs on this expanded set of features using 411 zebrafish larvae behaviour tracks (Fig. 2B). Fluoxetine, imipramine, LiCl and diazepam elicited strikingly similar behavioural syllables during the startle period (Fig. 2B). All AA drugs increased turning behaviour and decreased thigmotaxis distance and time, and spurting behaviours at lower doses. Only diazepam (at 1 µM or higher) diverged somewhat in that thigmotaxis distance was increased. Ketamine mimicked MK801 at high doses (100 µM) in that both increased thigmotaxis, whereas turning increased with 10 µM ketamine or 1 µM MK801, consistent with the effect being mediated by the NMDA receptor for which MK801 has higher affinity. Notably, increased turning was a prominent feature of the AA-drug treated fish (Fig. 2B). MK801 induced a large increase in motility at 10 µM consistent with the known hyperlocomotor effect of NMDAR inhibition in rodents and fish ^38,40–42^, however when measuring the startle phase only, the overall effect of ketamine on motility was minor, even though for stimuli it increased motility (Supplementary figure 6A, B). Interestingly, during the post-startle phase shown in figure 5, MK801 and ketamine responses were almost identical.

**Figure 2.**
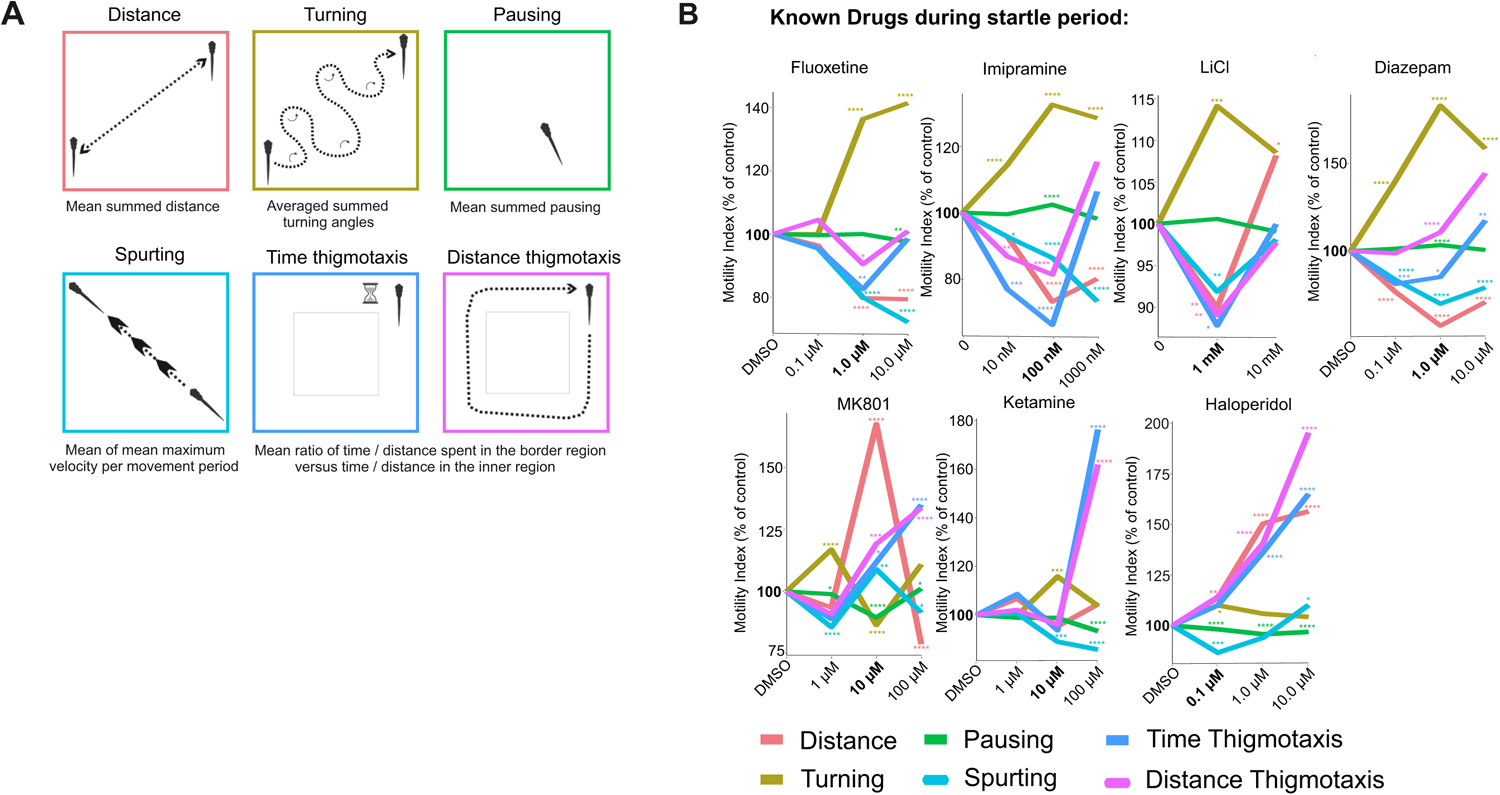
Testing the effect of AA drugs on zebrafish larvae behavioural sequalae during the 1 min startle phase. (A) The behavioural features extracted from zebrafish larvae tracking are shown. Distance, turning, pausing, spurting, time and distance thigmotaxis are extracted using R statistical computing platform. (B) Fish were exposed to AA drugs and ketamine, MK801 or Haloperidol as indicated and features (distance, turning, pausing, spurting, time and distance thigmotaxis) were extracted from the entire 1 min startle period. Averaged data indicates change relative to control for each of these behaviours according to color code. P-values were calculated by Wilcoxon Rank Sum test and adjusted with Benjamini-Hochberg procedure, where * p-value≤0.05; ** p-value *≤ 0.01; ***p-value≤0.001; ****p-value≤0.0001.

### Identification of JNK1 pathway signalling hubs

Having established phenotypic responses to clinically used AA drug classes in zebrafish larvae, we next turned our attention to the JNK1 pathway. As JNK1 regulates many physiological processes in brain, we were interested to identify downstream signalling hubs that could serve as alternative, possibly more specific AA drug targets than JNK1 itself. Initially we identified JNK1-regulated phosphoproteins by comparing the phosphoproteome from wild-type (WT) and *Jnk1-/-* mouse brains. Phosphoproteins that differed significantly by more than 1.5 fold in *Jnk1* knockout brain are presented as a circular array (Fig. 3A). We next predicted the most highly connected phosphoproteins to the JNK1-regulated ones using the GeneMANIA physical interaction network database. The predicted proteins that physically interact with the JNK1-regulated phosphoproteins are displayed at the centre of the circle (Fig. 3A). From the identified interacting proteins, we determined 29 candidate hubs from the JNK1-regulated phosphoproteome based on interaction counts and interaction confidence levels. To assess the significance of these interactions, we compared interaction counts for JNK1-regulated phosphoproteins to 1000 randomly generated datasets derived from the entire brain phosphoproteome (Fig. 3B). These hubs demonstrated significantly higher connectivity in *Jnk1-/-* mouse brains compared to WT brains. From among the hubs, we further selected those for which small molecule drugs were available for testing in the zebrafish behavioural screen. The selected hubs included YWHAZ/E (14-3-3s), AKT1, GSK3B, BRAF, and PKCE/G.

**Figure 3.**
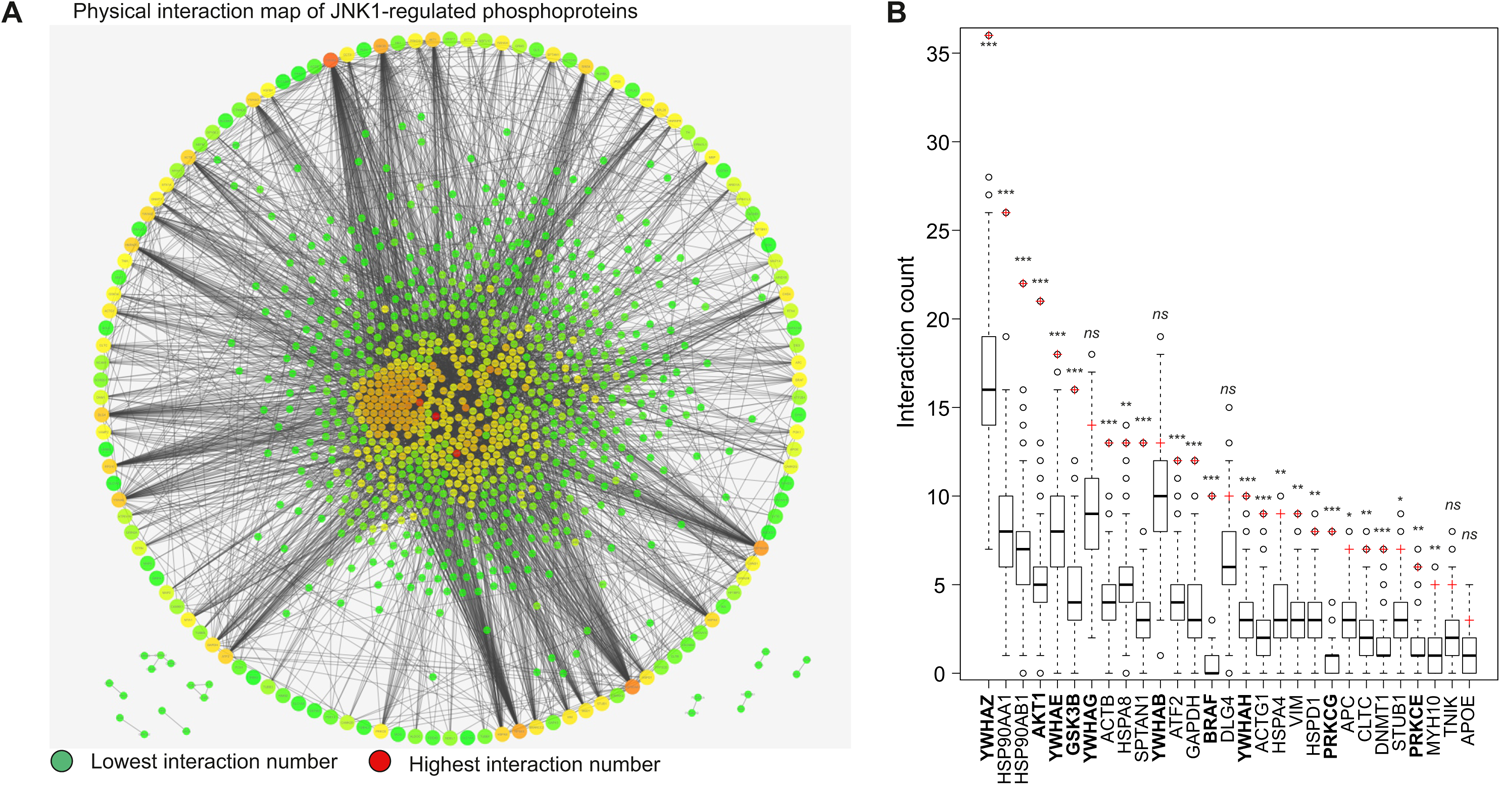
Identification of signalling hubs downstream of JNK1 in mouse brain. (A) JNK1 regulated phosphoproteins from *Jnk1-/-* mouse brain are organised in a large outer circle with connections to the predicted interacting proteins in the centre. Warmer colours indicate a larger number of physical interactions, red being the highest. (B) Signalling hubs derived from the *Jnk1-/-* mouse brain phosphoproteome (labelled using gene names) represent the most highly connected phosphoproteins from among all phosphoproteins that are significantly altered in *Jnk1-/-* brain verses wild-type. For each hub, the distribution of physical interaction count from the 1000 background networks is represented with a boxplot. The number of physical interactions for the same hub in the *Jnk1-/-* mouse brain phosphoproteome (ie the phosphoproteins that are significantly altered in *Jnk1-/-* mouse brain phosphoproteome), is depicted with a red cross. The most highly ranked JNK1-regulated signalling hubs are shown.

### JNK1 pathway hub compounds induce AA-like stereotypic behaviours during the startle period

We next screened JNK1 pathway drugs during the startle period. JNK inhibitors SP600125 at 1 µM, and JNK-IN-8 at 10 nM mimicked the AA drug profiles (Fig. 4A). Downstream of JNK1, we targeted signalling hubs 14-3-3 and AKT using R18, an inhibitor of 14-3-3 client binding ^43^ and SC79 (an AKT activator). This also induced a profile similar to that obtained with clinically used AA drugs. In contrast AKTi, an inhibitor of AKT showed an opposite effect on distance and pausing and less effect on turning (Fig. 4A). GSK-3 inhibitor (SB216763) at 1 µM and LiCl at 1 mM, both known to inhibit GSK-3, produced similar responses. We also screened PKC-targeted drugs. PKC-epsilon activator (FR236924)^44^ at 0.1 µM increased turning and pausing while decreasing distance (Fig. 4A). Bryostatin-1, a mixed PKC agonist, also increased turning, but differed from FR236924 in that it reduced spurting. Also at 10 nM (a dose that activates PKCα and PKCδ isoforms, but antagonizes PKCγ), it increased thigmotaxis and decreased pausing. Conversely, the pan-PKC inhibitor 3,4-Bis(3-indoyl)maleimide (BIM) increased turning and decreased thigmotaxis at 100 nM, a dose that inhibits PKCα more effectively than PKCε ^45^

**Figure 4.**
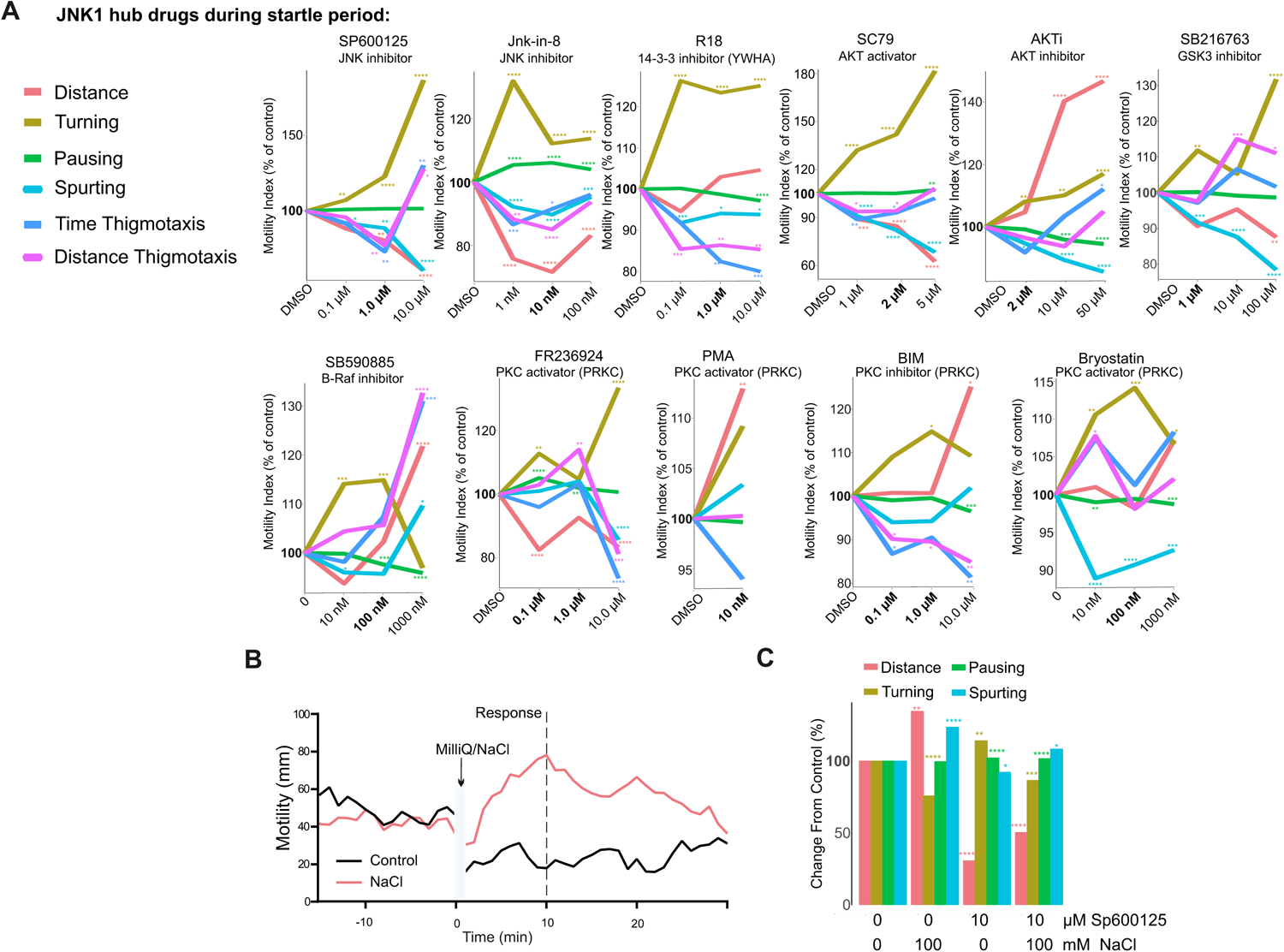
Testing the effect of JNK1 pathway hub drugs on zebrafish larvae behaviour during the startle period. (A) Zebrafish behaviours are shown during the 1 min startle period following treatment with pharmacological inhibitors or activators of JNK and downstream signalling hubs. As above, drug treatments were for 1 h before exposure to the startle battery. Mean distance, turning, pausing, spurting, time and distance thigmotaxis are shown for the following numbers of fish measurements (5 cycles per zebrafish larvae) JNK-IN-8: 346, SP600125: 224, Haloperidol: 220, SB216763: 358, FR236924: 360, R18: 354, SC79: 318, BIM: 216, PMA: 115, SB590885: 206, bryostatin-1: 359, AKTi: 360. P-values were calculated by Wilcoxon Rank Sum test and adjusted with Benjamini-Hochberg procedure and are indicated as follows * p-value≤0.05; ** p-value *≤ 0.01; ***p-value≤0.001; ****p-value≤0.0001. (B) Mean motility profiles of zebrafish larvae before and after milliQ (n=24) or NaCl (100 mM, n=24). (C) Mean distance, turning, pausing and spurting during the first 10 min following 100 mM NaCl was measured in the presence or absence of JNK inhibitor SP600125 (10 µM) or DMSO carrier in 7 dpf larvae. The number of fish per group were as follows: control = 47, NaCl + DMSO = 47, SP00125 + E3 = 48, SP00125 + NaCl = 45. Adjusted p-values are indicated as follows *p-value<0.05; ** p-value *< 0.01; ***p-value<0.005. n.s. = not significant. P-values were calculated using hypergeometric test and P-values were adjusted using the Benjamini-Hochberg procedure.

We also evaluated whether JNK regulated HPI axis stress in this model. To this end, we used saline stress which has been shown to elicit cortisol-dependent motility in zebrafish larvae that is HPI axis dependent ^46^. This treatment increased the distance moved and spurting, and decreased turning (Fig. 4B, C). Notably, pre-treatment with JNK inhibitor SP600125 (10 µM) for 1 hour significantly reduced these stress-induced behaviours (Fig. 4C). This identifies turning, pausing, spurting and distance as features that are JNK and HPI axis dependent from zebrafish motility tracking.

### Drug behaviour profiles during the post-startle period

The post-startle period is also relevant for stress responses and habituation ^47^, therefore we examined zebrafish behaviour during this period. Clinically validated drugs produced behaviours similar to those during the startle period but with a more pronounced reduction in distance moved, increased pausing, and milder effects on thigmotaxis (Fig. 5A). Turning behaviour during the post-startle response period increased more linearly with dose, and at 10 µM, ketamine’s behaviour profile resembled classical AA drugs in this period. Among the JNK1 pathway hub compounds, several induced anxiolytic-like profiles during the post-startle period (Fig. 5B). For example, the 14-3-3 inhibitor R18 (0.1 µM) mimicked classical AA drug profiles, reducing distance moved while increasing turning and pausing. The selective PKCε activator FR236924 (0.1 µM) showed a similar effect, decreasing distance and thigmotaxis while increasing turning and pausing.

**Figure 5.**
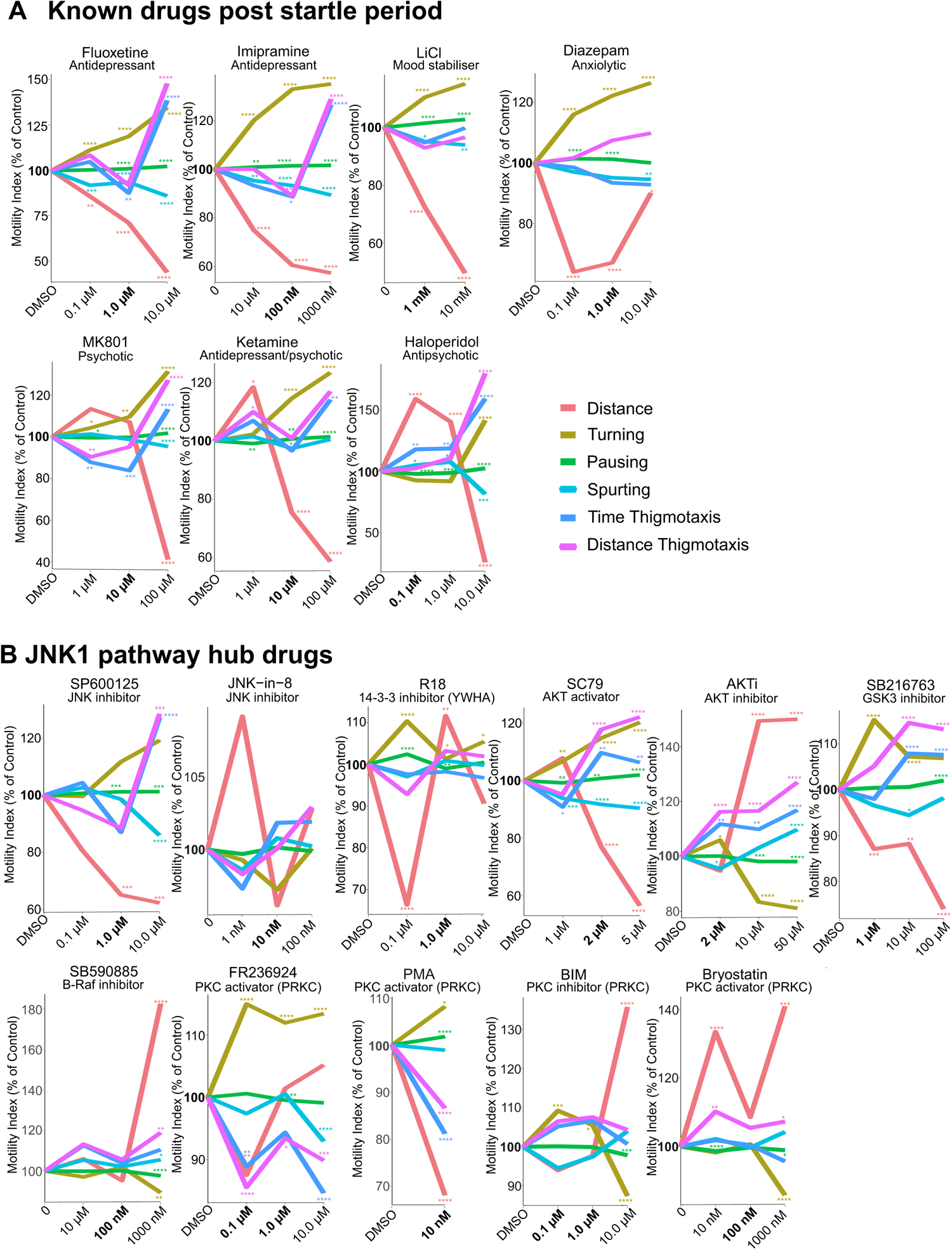
Testing the effect of AA drugs and JNK1 pathway hub drugs on zebrafish larvae behaviour during the post-startle period. (A) Fish were exposed to AA drugs and ketamine or MK801 doses as indicated and new features (distance, turning, pausing, spurting, time and distance thigmotaxis) were extracted from the first 10 min following the 1 min startle period. Averaged data from the following fish measurement numbers: diazepam: 337, fluoxetine: 338, imipramine: 335, LiCl: 210, ketamine: 325, MK801: 323 are shown. P-values were calculated by Wilcoxon Rank Sum test and adjusted with Benjamini-Hochberg procedure and are indicated as follows: * p-value≤0.05; ** p-value *≤ 0.01; ***p-value≤0.001; ****p-value≤0.0001. (B) Zebrafish behaviours during the 10 min following the 1 min startle period are shown with or without treatment with pharmacological inhibitors or activators of JNK and downstream signalling hubs. As above, drug treatments were for 2 h before exposure to the startle battery. Mean distance, turning, pausing, spurting, time and distance thigmotaxis are shown for the following numbers of fish measurements (from 5 cycles per zebrafish larva): JNK-IN-8: 346, SP600125: 224, Haloperidol: 220, SB216763: 358, FR236924: 360, R18: 354, SC79: 318, BIM: 216, PMA: 115, SB590885: 206, bryostatin-1: 359, AKTi: 360. P-values were calculated by Wilcoxon Rank Sum test and adjusted with Benjamini-Hochberg procedure and are indicated as follows * p-value≤0.05; ** p-value *≤ 0.01; ***p-value≤0.001; ****p-value≤0.0001.

JNK inhibitor SP600125 (1 µM) reduced distance, decreased thigmotaxis, and increased pausing, while the less well characterised JNK-IN-8 at 0.01 µM, had no significant effect. The AKT activator SC79 (2 µM) produced a response similar to JNK inhibitor and AA drugs, while the AKT inhibitor AKTi (50 µM) induced an opposite response, possibly representing anxiogenic-like behaviours.

GSK-3 inhibitor SB216763 mimicked the response of AKT activator SC79. The PKC activators FR236924 (0.1 µM) and PMA evoked similar responses, although Bryostatin-1, which activates some PKC isoforms, differed substantially in profile ^45^. Notably, NMDA receptor antagonists MK801 and high-dose ketamine showed markedly different dose response profiles during and after the startle period (Fig. 5), with post-startle responses aligning more closely with AA drugs. AKTi, BIM, and Bryostatin-1 did not fit the AA phenotype, as expected, whereas haloperidol fit both “AA” and “other” classification.

### Machine learning identifies compounds with anti-depressant/anxiolytic classification

In addition to traditional statistical analysis, we employed machine learning to create a comprehensive AA phenotype classification. For this we used Receiver Operating Characteristic (ROC) curves to assess the “fit” of test drugs to this classification. This approach has the advantage of overcoming the limitations and biases of regular statistical analysis and enhances the interpretation of complex data. We tested three supervised learning models: Random Forest classification-regression, GLMNET linear-regression, and SVM non-linear-regression. The SVM non-linear regression model achieved the highest training accuracy (>0.99) for both the startle and post-startle response periods with AA drugs, making it the chosen model to validate the test compounds.

Using motility tracking from the startle response period, we foud that SP600125, JNK-IN-8, SC79, GSK-3 inhibitor (SB216763), PKC isoform activators (FR236924 and PMA), and the 14-3-3 inhibitor (R18) classified as “AA” drug types with an area under the curve (AUC) of >97% in the SVM ROC analysis. For JNK inhibitors, this aligned with mouse data where JNK1 inhibition exerts an anxiolytic effect (Fig. 6A, B) ^29^. Conversely, inhibitors of the same molecules (BIM, AKTi, SB59085, and bryostatin-1) did not fit the “AA” category. Interestingly, Haloperidol, which is a D2 receptor antagonist used for its anti-psychotic drug effect, classified with both “AA” and “other” phenotypes.

When using post-startle response data, the accuracy of predictions for test drugs was generally higher. Thus, SC79, R18, SB216763, PMA, and FR236924 scored highly for an AA phenotype in the post-startle period, even surpassing the scores of JNK inhibitors, suggesting that greater functional specificity is achieved by targeting downstream of JNK1. A summary of the multiparametric zebrafish larvae behaviour test and the impact of signalling hubs downstream of JNK on the AA phenotype are outlined in Figure 6C.

**Figure 6.**
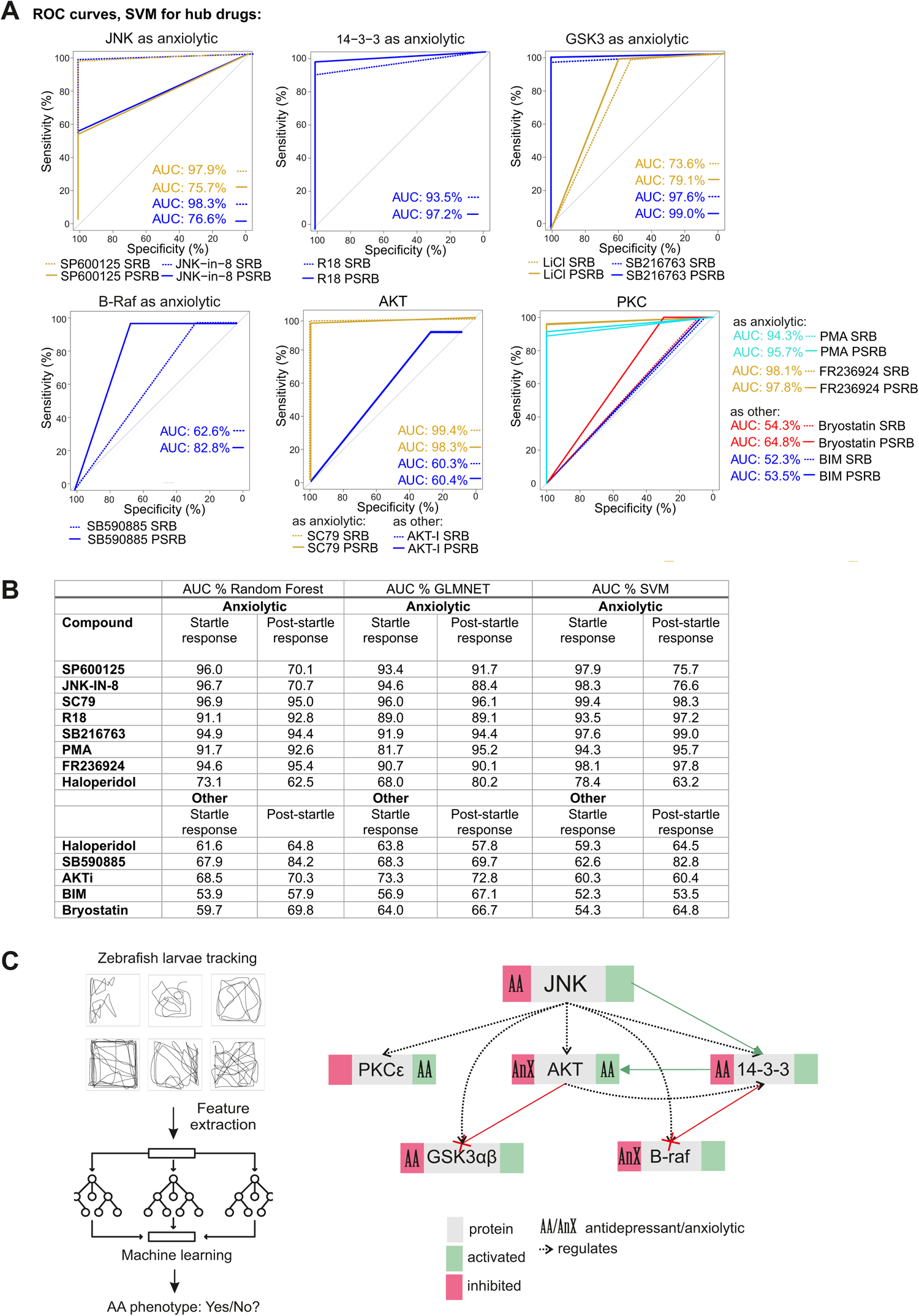
Machine learning predicts compounds with high sensitivity and specificity for generating an AA phenotype. (A) ROC curves show the Support Vector Machine (SVM) results for JNK pathway hub compounds to predict an “AA” or “other” phenotype during the Startle Response Behaviour (SRB) and the Post Startle Response Behaviour (PSRB). The number of fish measurements used for testing the machine learning models were JNK-IN-8: 346, SP600125: 224, Haloperidol: 220, SB216763: 358, FR236924: 360, R18: 354, SC79: 318, BIM: 216, PMA:, SB590885: 206, Bryostatin-1: 359, AKTi: 360. (B) A table summary of the area under the curve (AUC) values from Random Forest, Glmnet and SVM machine learning models are shown. Bold: Ranked highest overall taken from highest score across all machine learning approaches. The number of zebrafish tracks used for ML model training for the SRB class label “AA” were-diazepam: 350, fluoxetine: 335, imipramine: 343, LiCl: 210, and for class label “other” – ketamine: 349 and MK801: fish. During the post-startle response behaviour (PSRB) were as follows: class label “antidepressant or anxiolytic” – diazepam: 337, fluoxetine: 338, imipramine: 335, LiCl: 210; class label “other” – ketamine: 325 and MK801: 323. (C) A schematic summary of the zebrafish larvae behaviour screen for AA phenotype is shown. Beside it is a summary of results obtained using this test to evaluate the effect of JNK1 pathway hub drugs on AA-like behaviour.

## DISCUSSION

In this study, we establish a multi-parameter behavioural screen to classify a common AA phenotype in zebrafish larvae using clinically proven AA drugs. We use distinct groups of classical anti-depressant drugs; fluoxetine, representing a commonly used serotonin reuptake inhibitor; imipramine, a tricyclic anti-depressant, and ketamine, a fast-acting anti-depressant, as well as the mood stabilizing drug lithium chloride and diazepam, a commonly used anxiolytic of the benzodiazepine class. Despite having distinct pharmacological profiles, these drugs elicit similar effects on zebrafish larvae motility patterns. All AA drugs reduce distance moved, thigmotaxis and spurting, while they increase turning. This differs from the behavioural responses to psychosis-inducing drugs MK801 and high dose ketamine NMDA receptor antagonist drugs and anti-psychotic Haloperidol, and saline-induced HPI axis stress, all of which produce opposing behaviours in the same test battery. We employ machine learning to characterise an “AA phenotype” using all of these behavioural features, and go on to use the assay to evaluate the signalling hubs downstream of JNK1. The results uncover new regulators downstream of JNK1 that align with the classical AA-drug phenotype performing even better than JNK inhibitors, suggesting increased functional specificity.

Dysregulation of the hypothalamic-pituitary-adrenal axis leading to hypercortisolemia is one of the main circuit irregularities contributing to anxiety disorders and major depression ^48–51^. This hormonal change can trigger a cascade of events including the activation of JNKs. Accordingly, JNK is activated by cortisol and induces pro-inflammatory cytokines as well as amygdaloid dendritic hypertrophy ^31,52–54^. Additionally, JNK plays a role in the homeostatic regulation of glucocorticoid receptor transcriptional activity ^25,28^. In contrast, when JNK is inhibited in mouse brain, a reduction in anxiety- and depressive-like behaviours is observed ^55^, as is the case with JNK inhibitor treatment or genetic deletion ^29,56–58^. In contrast, chimeric JNK activation induces depressive-like behaviour and impaired assessment of risk/reward benefit ^58,59^. In zebrafish brain at the larval stage, JNK1 orthologs (*mapk8a* and *mapk8b)* are expressed ^60^, and consistent with behavioural findings in rodents ^54^, we find that treatment with structurally independent JNK1 inhibitors SP600125 and JNK-IN-8, induce an AA phenotype with A.U.C.s of 97.9% and 98.3% respectively in the startle period. Thus, JNK inhibitors elicit a behavioural response in zebrafish larvae that matches closely to that of clinically used AA drugs.

A main goal of this study was to identify signalling hubs downstream of JNK1 influencing anxiety- and depressive-like behaviours. We discovered 14-3-3ζ (YWAZ) and 14-3-3ε (YWHAE), members of the 14-3-3 client binding family ^61^, as key hubs in this process. JNK is known to phosphorylate 14-3-3ζ leading to client protein release ^62^. Consistent with this, the interaction count for ζ and ε 14-3-3 isoforms increases in *Jnk1-/-* mouse brain, indicating that JNK1 regulates 14-3-3 binding dynamics in the CNS. In zebrafish larvae, we show that R18 peptide which disrupts 14-3-3 client binding ^63^, recapitulates the AA-drug phenotype with A.U.C. of 93.5% and 97.2% in the startle and post-startle period respectively, consistent with the involvement of ζ or ε isoforms ^61,64^. 14-3-3ε has also been shown to mediate chronic stress-induced depression ^65^. Interestingly, fluoxetine rescues the behavioural deficit in zebrafish lacking 14-3-3ζ suggesting a shared mechanism with fluoxetine ^66^.

We also identify AKT as a potential signalling hub downstream of JNK1 based on mouse brain phosphoproteome interaction network analysis. AKT and 14-3-3 are mechanistically linked, thus inhibition of 14-3-3 client binding activates AKT ^67,68^, potentially placing AKT regulation by JNK1 downstream of 14-3-3. We found that the AKT activator SC79 evoked an AA phenotype with A.U.C.s of 99.4% and 98.3% for the startle and post-startle response periods in the zebrafish larva screen. Conversely, the allosteric inhibitor of AKT (AKTi) induced an opposite response (A.U.C. 60.3% and 60.4% respectively). That AKT activation induces an AA phenotype fits with the well-documented anti-depressant action of AKT in animal models and depressed subjects, and its regulation of BDNF ^69,70^. Consistent with this, we also identify that GSK-3β is a hub downstream of JNK1 in mouse brain. This is not surprising as GSK-3α/β isoforms are phosphorylated directly by AKT on S21 and S9 (α and β isoforms respectively), leading to GSK-3 inhibition ^71^. In the zebrafish larva assay, the GSK-3α/β inhibitor SB216763, elicits an AA phenotype with A.U.C.s of 97.6% and 99.0% during the startle and post-startle response phases.

B-Raf was another signalling hub identified downstream of JNK1. In the interaction screen, B-Raf displayed an increase in protein interactions (from “0” to “10”) in the phosphoproteome from *Jnk1-/-* mouse brain compared to wild-type. However, when we tested the B-Raf inhibitor SB590885, the resulting phenotype did not classify as “AA” either during the startle or post-startle period. Interestingly, B-Raf is also known to interact with 14-3-3, which functions to maintain its kinase domain in an inactive state ^72^. The expectation therefore would be that activation of B-Raf upon disruption of 14-3-3 binding may allow it to contribute to an anxiolytic-like state, as observed with the 14-3-3 client binding inhibitor R18. However, it was not possible for us to test B-Raf further due to lack of activators for this molecule. The oncogenic nature of B-Raf activating mutations contraindicates the use of such molecules.

Finally, we tested PKC activators and inhibitors in the zebrafish screen to explore their impact as possible effector hubs downstream of JNK1. We used the PKCε-activator FR236924 and the pan-PKC activator PMA, both of which evoked a clear anxiolytic-like effect (A.U.C.s of 97.8% and 95.7 % respectively). Consistent with this, the broad specificity PKC inhibitor (BIM) had the opposite effect. Notably however, in some contrast with our findings in zebrafish larvae, BIM treatment attenuates corticotrophin releasing factor (CRF) facilitation of the acoustic startle response in mice, although it had no effect in the absence of CRF ^73^. Moreover, *PKCγ*-/- mice exhibit low anxiety ^74^. Interestingly also, that PKCε activator (FR236924) promotes an AA phenotype in zebrafish is supported by the response to byrostatin-1 at 100 nM, (a dose that activates PKCε ^75^) during the startle response, even though when analysed at the full dose response in our study the overall phenotype is “other”. While these studies suggest isoform-specific roles for PKC variants in anxiety-like behaviours, we identify that PKCε represents a signalling hub downstream from JNK1 which controls an AA phenotype in zebrafish larvae, indicating that further analysis of this isoform in the context of stress-related behaviours is warranted.

The observation of overlapping behavioural phenotypes elicited by drugs with distinct pharmacologies merits discussion. The primary target of fluoxetine is to block serotonin reuptake, and that of imipramine is to block serotonin and norepinephrine reuptake, and to additionally antagonise dopamine D2 and acetylcholine receptors ^76^. These drugs take weeks to elicit a notable effect on neurotransmitter levels in both responders and non-responders and other effects such as neuroplasticity and neurogenesis changes may be additionally needed for the therapeutic effect ^76,77^. A recent study identified that the Tropomyosin receptor kinase (TrkB) is a common binding site for a range of anti-depressant drugs including fluoxetine, imipramine and ketamine. The authors demonstrated that binding of these drugs to TrkB occurs within minutes and is required for the anti-depressant-like effect on behaviour ^78^. The authors rationalized that the well studied neuroplasticity effects of BDNF, mediated via TrkB binding, could explain the anti-depressant effect and its timeline. TrkB could represent the common mediator of drug responses measured here as zebrafish larvae indeed express TrkB orthologs (*trkB1* and *trkB2*) from 6 days post fertilization onwards ^79^. Moreover, activation of TrkB leads within minutes to the activation of AKT ^69^. We uncover that JNK1 signalling hubs converge on the AKT pathway with 14-3-3ζ/ε and GSK-3 in addition to AKT itself identified. Moreover, treatment of zebrafish larvae with the AKT activator (SC79) evokes an AA phenotype with closes match to the clinically used AA drugs with an A.U.C. of 99.4 %. AKT pathway activation is the most likely explanation for the temporal and phenotypic overlap observed in response to AA and JNK1 hub drugs.

Animal models are limited when it comes to modeling complex disorders such as anxiety and depression. They do not account for subjective experience, or complex genetic and environmental background, and the models that do exist in mice require large numbers to attain statistical power ^80,81^. Here we present an assay that is more straightforward and scalable than rodent models. It measures stereotypic responses to acoustic and light stress, based on larval behavioural sequalae (distance, turning, pausing, spurting, thigomataxis distance, thigmotaxis time, dose and size effect). This may lack face and construct validity, and it is not designed to model a specific disease mechanism. Nonetheless, it does measure HPI axis stress responses and the neuroendocrine and monoamine control systems underlying the stress axis is conserved from chordates to mammals ^82^, moreover locomotor activity in zebrafish larvae is inherently cortisol dependent ^12^. A similar approach has been used to identify novel anti-psychotic compounds in zebrafish larvae ^15^, and to investigate the paradoxical excitatory effects of GABA and serotonin ^16^, for example. Interestingly, here we also observe a similar hyperactivity effect with the GABAR agonist diazepam when fish are exposed to violet light and acoustic stimuli together (Fig. 1H). Our study in addition exploites machine learning to classify the desired phenotype, enabling use of complex combinations of features. The result is high accuracy prediction scores for test drugs that are known to be anxiolytic/anti-depressant in mice e.g. AKT activator and GSK-3 inhibitor ^69,70,83^. This highlights the potential of the zebrafish larvae screen to potentially identify compounds that mimic AA drugs, for further testing in rodent models, which is rather the goal of a phenotypic screen.

A possible explanation for the high prediction score could be linked to the dependence of the zebrafish larvae locomotor response on the HPI-axis ^46^, and the drugs used regulate the HPA-axis ^84^. Although the stereotypic behaviours measured here may not directly relate to AA conditions, JNK is activated by cortisol ^30,85^ leading to synapse withdrawal ^31^, which may influence circuit behaviour and long-term mental state. Notably also, the phenotype of AA-drugs differ from those of psychotic compounds MK801 and ketamine. Similarly, the anti-psychotic D2 antagonist Haloperidol shows dual classification which is consistent with its known paradoxical side effect, whereby some patients can respond with worsening of symptoms such as increased agitation, insomnia and hallucinations in patients ^12^. Consistent with this in the zebrafish larvae, we find that Haloperidol shows a distinct hyper-motility phenotype, which possibly accounts for its classification as “other”; yet it also increases turning and diminishes spurting, consistent with the “AA” phenotype (Fig. 2; Supplementary figure 7). This paradoxical effect to Haloperidol was also previously reported in zebrafish larvae ^15^. A long-term advantage of the screen is the potential reduction in the number of mice needed for drug discovery, while expanding the utility of zebrafish larvae in neuropsychopharmacology studies.

## MATERIALS AND METHODS

### Zebrafish husbandry

WT zebrafish (*D. Rerio*) larvae were derived from a cross between the high-performance AB strain and the Tu short-fin wild type strain ^20^. Fish were maintained in the Turku Bioscience Zebrafish Core facility at 28.5°C, under a 12 h day/night cycle. Embryos were cultured in E3 medium (5 mM NaCl, 0.17 mM KCl, 0.33 mM CaCl_2_, 0.33 mM MgSO_4_) with 30 eggs per 10 cm diameter dish. Larvae of indeterminate sex were used from 3 to 8 dpf as indicated. Experiments were carried out in the afternoon to maximise consistency of locomotor behaviour ^86^. Fish were bred in two tanks. At least two plates of fish were tested for every drug. There were minor differences in baseline motilities between plates therefore data was normalised to internal control fish measured on the same plate. Afterward the experiment, zebrafish larvae are anesthetized with 200mg/L Tricaine for 5min, and then euthanized by transferring to 1% chloramine ice-water solution. Behavioural experiments were carried out following the guidelines that were approved by the National Animal Experiment Board (ELLA) and the Regional State Administrative Office of Southern Finland (ESAVI) under licenses ESAVI/16343/2019 and ESAVI/30686/2022, which is the authority in Finland that ensures that animal experimentation is carried out following the law and thus follows EU ethical guidelines for animal experimentation. All all methods are performed and reported in accordance with the ARRIVE guidelines (https://arriveguidelines.org).

### Behavioural battery test

7 days post fertilization (dpf) zebrafish larvae were transferred to square well 96-well plates and drug dosages were administered following which the plate was directly transferred to the DanioVision^TM^ device (Noldus Information Technology, BV, Wageningen, Netherlands) and acclimatized for 1 h in the dark at 28.5 °C. Larvae then underwent the test battery consisting of 5 cycles of a 60 s string of acoustic/vibratory (tapping) and light stimuli programmed using Ethovision13 XT software (Noldus Information Technology, BV, Wageningen, Netherlands). The 60 s stimuli were as follows; high (H) and low (L) intensity patterns (1.6 & 4.2 watt): 1 s pause, 6x L-tap (0.5 s break between individual taps), 4 s pause, 4x H-tap, 1 s pause, 8x L-tap, 8 s pause, 2x L-tap, 10x H-tap, 7 s pause, 2x H-tap, 13 s pause, 14x H-tap, 3 s pause. Visual stimuli evoked using red (635 nm, 192 Lm), blue (470 nm, 240 Lm), purple (red and blue together) and white LEDs, switched colour at 10 s intervals in the following order: darkness, red, blue, purple (red & blue), white, and sequential flickering light of all colours sequentially at 0.1 to 0.2 s intervals. After the stimuli period there was a 29-minute intermission period. This 30 min sequence was repeated 4 more times. Digital video tracking was performed using the Daniovision imaging unit using a pre-programmed protocol with the included Ethovision XT software. Acquisition was 60 fps and exported as csv file for automated analysis using customized software in R. The data analyst was agnostic to the drug properties.

### Stress response assay

5 or 7 dpf larvae (as indicated) were transferred to a 48-well plate with 1 fish/well and acclimatized for 1 h following which they were exposed to stress: either with white light illumination (10,000 Lux) for 50 s or addition of 100 mM NaCl. Larvae were tracked for a further 30 min following stressor in the dark using infrared light. NaCl experiments were performed using ambient level (500 Lux) white light.

### Drug treatments

Larvae at the indicated ages were treated with increasing doses of drugs and compounds as follows: fluoxetine, diazepam, FR236924 and PMA (Phorbol 12-myristate 13-acetate) and ketamine were from Tocris Bioscience, Abingdon, UK; haloperidol, imipramine, LiCl, SP600125, SB590885, Bryostatin-1 and MK801 from MERCK Sigma Aldrich Solutions, Darmstadt, DE; R18 from Enzo Life Sciences distributed by AH Diagnostics, Helsinki, FI; SB216763, JNK-in-8 and AKTi from Selleck Chemicals LLC, Houston, Texas; bisindolylmaleimide I (BIM) from Santa Cruz Biotechnology Inc., Dallas, Texas and SC79 from R&D systems, McKinley Place NE, Minneapolis or NaCl in E3 medium. Treatments were for 1h at 28.5 °C before commencing the startle battery. Control fish were treated with equivalent volume of carrier.

### Behaviour data analysis – Motility

Raw motility tracking data was exported in csv format, where each file contained data from 1 experimental cycle, and each worksheet contained time series data from 1 fish. Relative motility index (M.I._R_) (plots of left side) represents the motility during a certain condition (MC_x_) relative to the motility during the control condition (MC_0_). The relative motility index was calculated as follows from all 5 cycles:

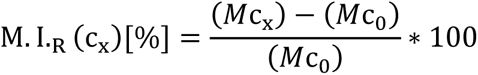

“Relative Peak Analysis” (shown in Fig. 2 right side panels) represents the motility during peaks P1, P2, P3, P4, P5, P6 of a test condition relative to the same peaks from control fish in the same plate. The data scientist handling the analysis was agnostic to the drug treatments. Analysis was done in R.

### Analysis of other features

X and Y positions, heading, and distance features from the time series data were utilized to calculate behavioural features: distance, turning, pausing, spurt velocity, and thigmotaxis by distance and time.

#### Distance

Distance was summed from all time points per fish and mean total distance travelled per treatment dose was calculated.

#### Turning

Turning angles between consecutive time points were calculated from the heading parameter of the last and current time points, and converted to magnitude representing the smaller angle in 360 degrees. Relative turning angles were calculated by summing all time points for each fish, then normalizing to distance travelled, and presented as average values according to drug dose.

#### Pausing

Time points with value 0 for distance were summed to give a value for total pause time for each fish.

#### Spurt velocity

Time bouts between pauses were identified and the velocity for each time point was calculated from distance feature. The maximum velocity from each time bout was recorded and the mean maximum velocities from all time bouts in a single fish were calculated.

#### Thigmotaxis

The well border coordinates from the 96-well plate were recorded from Ethovision graphic user interface (GUI). An inner border was (virtually) created that was 40% of the width and length of the original border, and placed in the centre of each well. The coordinates of these inner borders served as thresholds for thigmotactic behaviour. Time points where a fish travelled outside the inner border were identified and the ratio of near outer border travel versus total travel for each fish was calculated for time and distance. Anaysis was done in R.

### Machine learning analysis

Machine learning algorithms were applied to zebrafish larvae behaviour data during the 1 min startle period and during the 10 min of the recovery period. Specifically, the quantitative variables total pause duration, total distance traveled, relative turning angle, average spurt speed, total thigmotaxis time and total thigmotaxis distance were normalized against control (or zero-dosage fish) data as percentages. These normalized metrics taking into account effect sizes, along with dosage level variable, were incorporated as features for the machine learning models. The models were tasked to differentiate between “antidepressant or anxiolytic” drugs and “other” drugs. For the training dataset, drug treatments including diazepam, fluoxetine, imipramine, and lithium chloride (LiCl) were categorized as “antidepressant/anxiolytic” class, while ketamine and MK801 treatments were categorized as “other”. Generalized Linear Model via penalized maximum likelihood (Glmnet), Support Vector Machine (SVM) with a polynomial kernel, and Random Forest algorithms were utilized to build 3 separate models each for startle response behaviour (SRB) data during the 1 min startle battery and post-startle response behaviour (PSRB) data during the 10 min recovery period. The constructed models were subsequently employed to predict the class category of other drug treatments. Machine learning was performed with “caret” package within the R environment.

### Statistics

All behavioural features were compared to control fish from the same experiment. For statistical testing, feature values were collected into a distribution, and this data according to drug dose was compared to respective control distributions using Wilcoxon signed-ranked test or one-way or two-way ANOVA with Tukey or Dunnett’s multiple comparison test as indicated. Calculations and plots were made using a customized pipeline in R programming language. Data analysis was automated and the data scientist was agnostic to the treatment drugs.

### Preparation of samples for mass spectrometry, SDS-PAGE and in-gel digestion

All the chemicals for digestion and mass spectrometry analysis were from Sigma-Aldrich (Stockholm, Sweden). Whole brain was isolated from WT and *Jnk1-/-* C57B6J mice ^87^ at embryonic day 15 (E15), post-natal day 0 (P0), post-natal day 21 (P21), and 8 months (Adult). We extracted brains from three mice for each age and genotype and snap froze them in liquid N_2_ and then pulverized them in a Micro-dismembrator II (Braun Biotech International, Melsungen, Germany). We weighed the powder and stored aliquots at −80°C before use. SDS (1 %), supplemented with protease and phosphatase inhibitors (Sigma-Aldrich, St. Louis, USA), was added to the brain-powder. Mechanical disruption followed using a syringe twice. We centrifuged samples for 1 hour at 4°C and collected the supernatant. We quantified protein concentration using the Total Protein Kit, Micro Lowry, Peterson’s Modification (Sigma-Aldrich, St. Louis, USA).

A total protein amount of 100 μg for each sample was separated on 12% Criterion gels (Bio-Rad Laboratories, Hercules, Ca, USA), following the manufacturer’s instructions. We washed the gel in milliQ, stained for 30 min with GelCode (Bio-Rad Laboratories) and washed overnight in milliQ before manually slicing each lane into 5 equal slices. We de-stained gel slices by washing 3 times in 25 mM ammonium bicarbonate (AMBIC)/50% acetonitrile and dried in a vacuum centrifuge (Speedvac). Samples were reduced (10mM DTT/100 mM AMBIC, 1 h at 56°C), alkylated (55 mM iodoacetamide (IAA)/100 mM AMBIC, 45 min at room temperature in the dark), washed 2 times with 100 mM AMBIC and dehydrated with ACN before being dried in the Speedvac. We rehydrated slices with 12.5 μg/ml modified porcine trypsin (Promega, Madison, WI, USA) in 50 mM AMBIC and digested overnight at 37°C. Peptides were eluted 2 times in 75% ACN/5% FA, dried 1 h in the Speedvac and dissolved in 0.1% formic acid.

### TiO_2_ phospho-peptide enrichment

we incubated samples for 10 min with Buffer A (1 M GA, 80% ACN, 5% TFA) followed by incubation with the TiO2 Mag Sepharose (GE Healthcare Life Sciences) for 30 min in an Eppendorf Thermomixer at 800 rpm. Liquid was removed and washing steps were done using Buffer A and 2 times using Buffer B (80% ACN, 1% TFA). Phospho-peptides enriched were eluted 2 times using 100 mM ammonium hydroxide in water and dried 1h in the Speedvac.

### LC-MS/MS analysis

Dried peptides were resuspended in 0.1% formic acid and separated using an Eksigent nano-LC 2D system (Eksigent, Dublin, CA, USA) consisting of a solvent degasser, a nano-flow pump and a cooled auto-sampler. Peptide concentrations were measured by Nanodrop (ThermoFisher, Stockholm) and the concentration adjusted to the same amount in all samples. Eight μl of sample was loaded and washed for 15 min onto a pre-column (Acclaim PepMap 100, C18, 3 μm particle size, 50 μm diameter, Thermo Fisher Scientific, Hägersten, Sweden) at a constant flow of 5 μl/min solvent B (0.1% FA in ACN). We ran three biological replicates and two technical repeats for each of the two genotypes at the four time points. Phosphopeptides were loaded into a RP analytical column (10 μm fused silica emitter, 75 μm ID column, 16 cm Pico Tip, New Objective) packed in-house with C18 material ReproSil-Pur and separated using an eighty-minute gradient from 3% to 40% solvent B at a flow rate of 300 nl/min. The gradient was followed by 20 min column washing with 90% ACN, 0.1% FA and 15 min re-equilibration with 3% solvent B. The HPLC system coupled to an LTQ-Orbitrap XL mass spectrometer (ThermoFisher Scientific, Bremen, Germany) operated in Data Dependent Acquisition mode. Spray voltage was set to 1.90 kV and the temperature of the heated capillary was set to 200 °C. The ten most intense ions from the survey scan performed by the Orbitrap were fragmented by collision-induced dissociation (CID) in the LTQ (normalized collision energy 35, parent mass selection window 0.5 Da, 30 ms activation time and minimum signal threshold for MS/MS scans set to 100). We excluded unassigned charge states and singly charged ions from fragmentation. The dynamic exclusion list was limited to a maximum of 500 masses with a retention time window of 2 minutes and a relative mass window of 10 ppm. The X-calibur software version 2.0.7 (Thermo Scientific) controlled the HPLC, mass spectrometer and data acquisition.

### Shotgun data analysis from LTQ Orbitrap

The raw data was uploaded into Progenesis LC-MS (Waters) for analysis. After raw mass spectra were aligned and normalisation across all runs was carried out by total ion current in the analysis area manually defined for the reference HPLC for the time and mass ranges. We submitted the raw files from the mass spectrometer to Thermo Proteome Discoverer (version 1.0.43.0) for protein identification by Mascot. Cysteine carbamidomethylation was set as a fixed modification, methionine oxidation and phosphorylation of serine, threonine, tyrosine (STY) were set as variable modifications. Data were searched against Uniprot *Mus musculus* database containing 170.598 sequences (release date April 2014). The mass tolerance was set to 10 ppm for the precursor ion and 0.8 Da for the fragment ions. Trypsin was set as the protease and a maximum 2 of missed cleavages were allowed. We ran raw data using an FDR of 5% and a score cutoff of 20. We discarded proteins marked as contaminants, reverse hits and “non-unique peptides”. We selected phospho-peptides with a Mascot score higher than 20 and verified using the default Mascot Delta score of 0.1 for comparative analysis between groups.

### Data merging and statistical analysis phosphoproteomics data

Downstream analysis of MS intensity outputs from Progenesis LC-MS (Waters) utilized a customized R (v.3.4.0) software pipeline. Phosphopeptide data were refined as follows: (i) all phosphopeptides from 5 gel slices were merged, (ii) identical phosphopeptides, i.e. those with identical sequence, UniProt identifier and number and position of phosphorylated residues, were merged by summing the intensities. (iii) If a peptide entry had >1 missing value across 3 replicates, all entries were removed. For quality control of data, a distribution plot of phosphopeptide intensities for each unique genotype at given ages was visualized using boxplot and histogram analysis, using ggplot2 ^88^ (Supplementary F1). “Phosphopeptide-centric” and phosphosite-centric analysis was performed as appropriate and is defined in figure legends. We used the Bioconductor package RankProd ^89,90^ for statistical analysis. We calculated mean ratios of phosphopeptide intensities [*Jnk1-/-/*WT] per entry (for both phosphopeptide-centric and protein-centric changes). Only phosphorylation intensity changes that passed a threshold Ratio*_Jnk1-/-_*_/WT_ > 2, with p-value <0.05 were included.

### Hub analysis

29 candidate hubs (Fig. 2H, I) that were significantly altered in *Jnk1-/-* mouse brain compared to wildtype brain and were part of the psychiatric disease associated genes according to the MetaCore^TM^ database schizophrenia gene list were selected to analyse as potential signalling hubs acting downstream of JNK1 in regulating depressive/anxiety-like behaviour. The selection criteria included a threshold for phosphoproteins that showed either ≥ 7 physical interactions, or > 3 high weight (weight > 0.04) physical interactions from the physical interaction network built with GeneMANIA from among the significantly altered phosphoproteins derived from the wildtype and *Jnk1-/-* mouse brain phosphoproteomes. 29 proteins fulfilled these criteria. To control for hub protein’s inherent functional plasticity variations, 1000 control phosphoprotein networks of the same size as the *Jnk1-/-* psychiatric disease phosphoprotein network were randomly generated from the entire brain phosphoproteome dataset using GeneMANIA (v.3.4.1). The number of interactions of the 29 proteins from the significantly altered phosphoproteome network in *Jnk1-/-* mouse brain was each compared to the distribution of interaction count from 1000 randomly generated control networks to determine the significance of the hubs. Statistical significance for the hubs was calculated using the hypergeometric test. P-values were adjusted using the Benjamini-Hochberg procedure.

### Receiver Operating Characteristic (ROC) analysis

ROC curves were constructed to visualise the predictive proficiency of the machine learning models. Individual curves were plotted for each test drug from the test dataset, paired with a control drug that was either classified as a “positive control” or a “negative control”. Each treatment drug is assigned with a “true” class label that reflected its known properties and molecular interactions. Drugs of the “antidepressant or anxiolytic” class were coupled with the “negative control” drug ketamine, which provided the “true” class label of “other”. Conversely, drugs classified under the “other” class were paired with the “positive control” drug fluoxetine for the “true” class label of “Antidepressant or anxiolytic” for the purposes of ROC curve calculation. The construction of the ROC curve was facilitated by the ‘pROC’ package within the R statistical programming environment.

### Data availability

The phosphoproteomics data generated during the current study is available in the PRIDE repository, (www.ebi.ac.uk/pride/).

## AUTHOR CONTRIBUTIONS

B.H., C.S. J.P. and J.J. carried out experimental work and data analysis. Y.H. carried out the phosphoproteomic and bioinformatics data analysis, machine learning. C.S., Y.H. and E.C. wrote the manuscript. All authors contributed to the manuscript and figure preparation. I.P. supervised the fish husbandry and contributed to data interpretation. E.C. conceptualized and supervised the project.

## ACKNOWLEDGEMENTS

We thank Zebrafish Core (Turku Bioscience Centre, University of Turku and Åbo Akademi University) supported by Biocenter Finland for assistance.

