## Supplementary Materials for "JNK1 And Downstream Signalling Hubs Regulate Anxiety-Like Behaviours In A Zebrafish Larvae Phenotypic Screen"

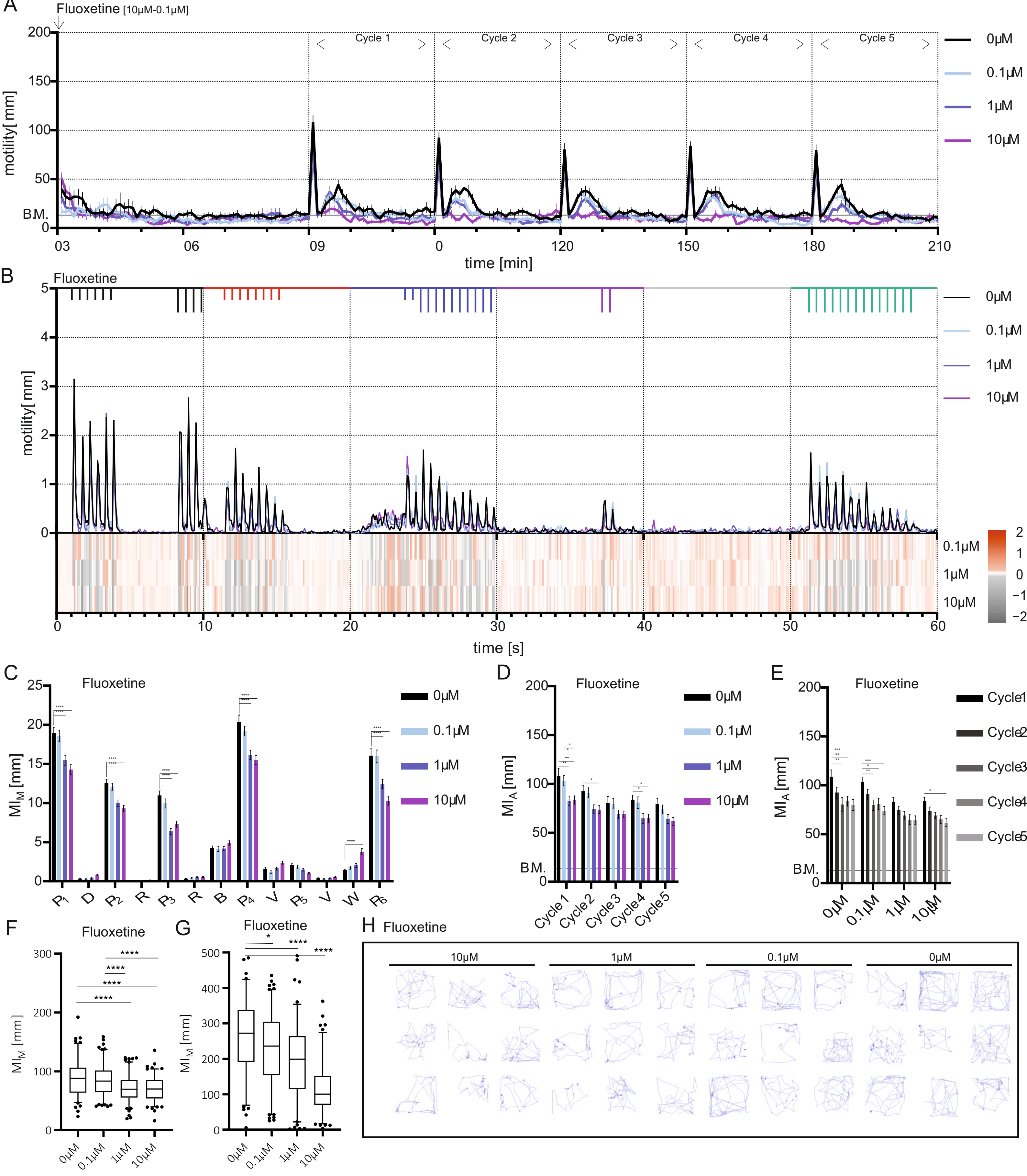


Supplementary figure 1 - Fluoxetine

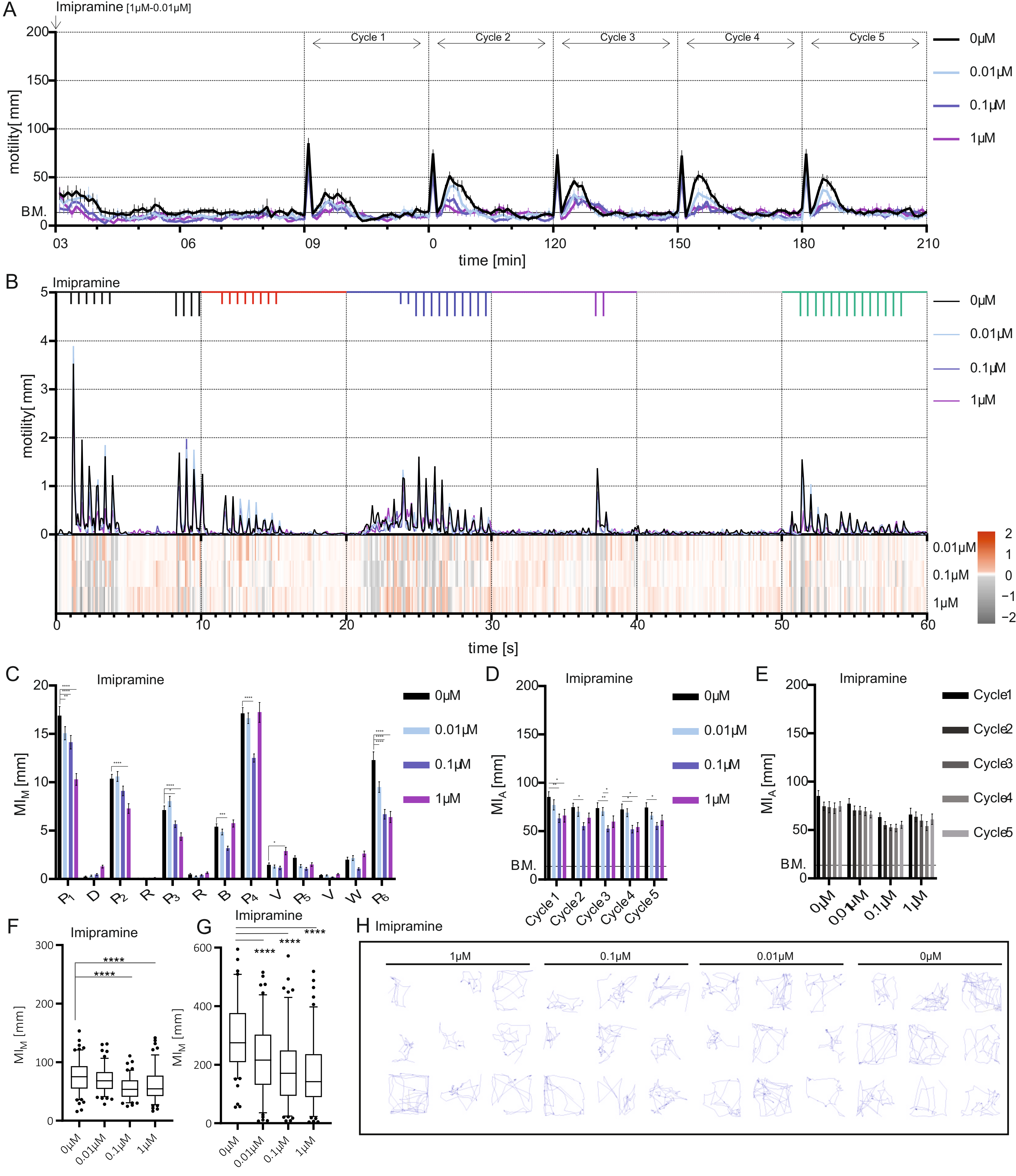


Supplementary figure 2 - Imipramine

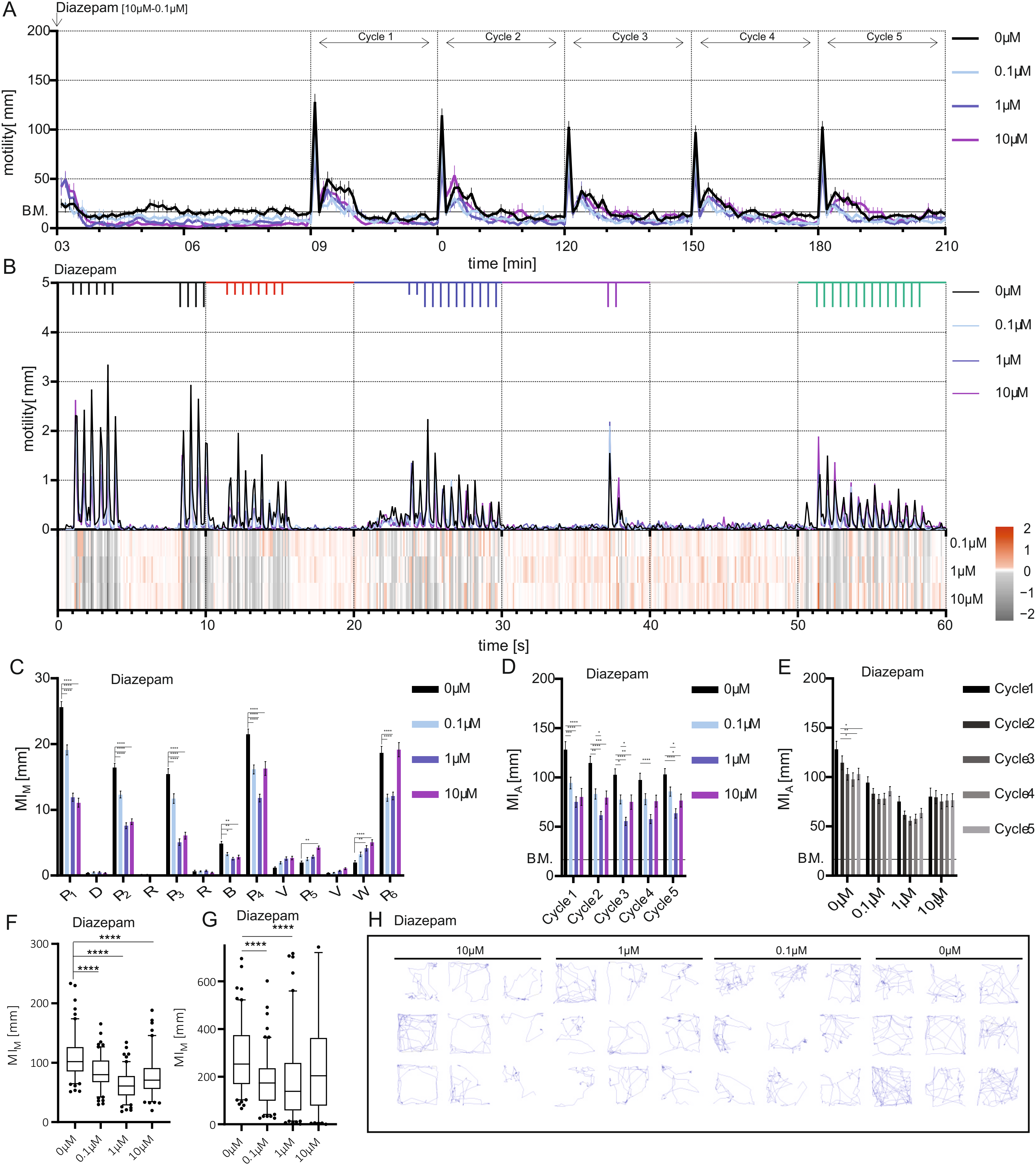


Supplementary figure 3 - Diazepam

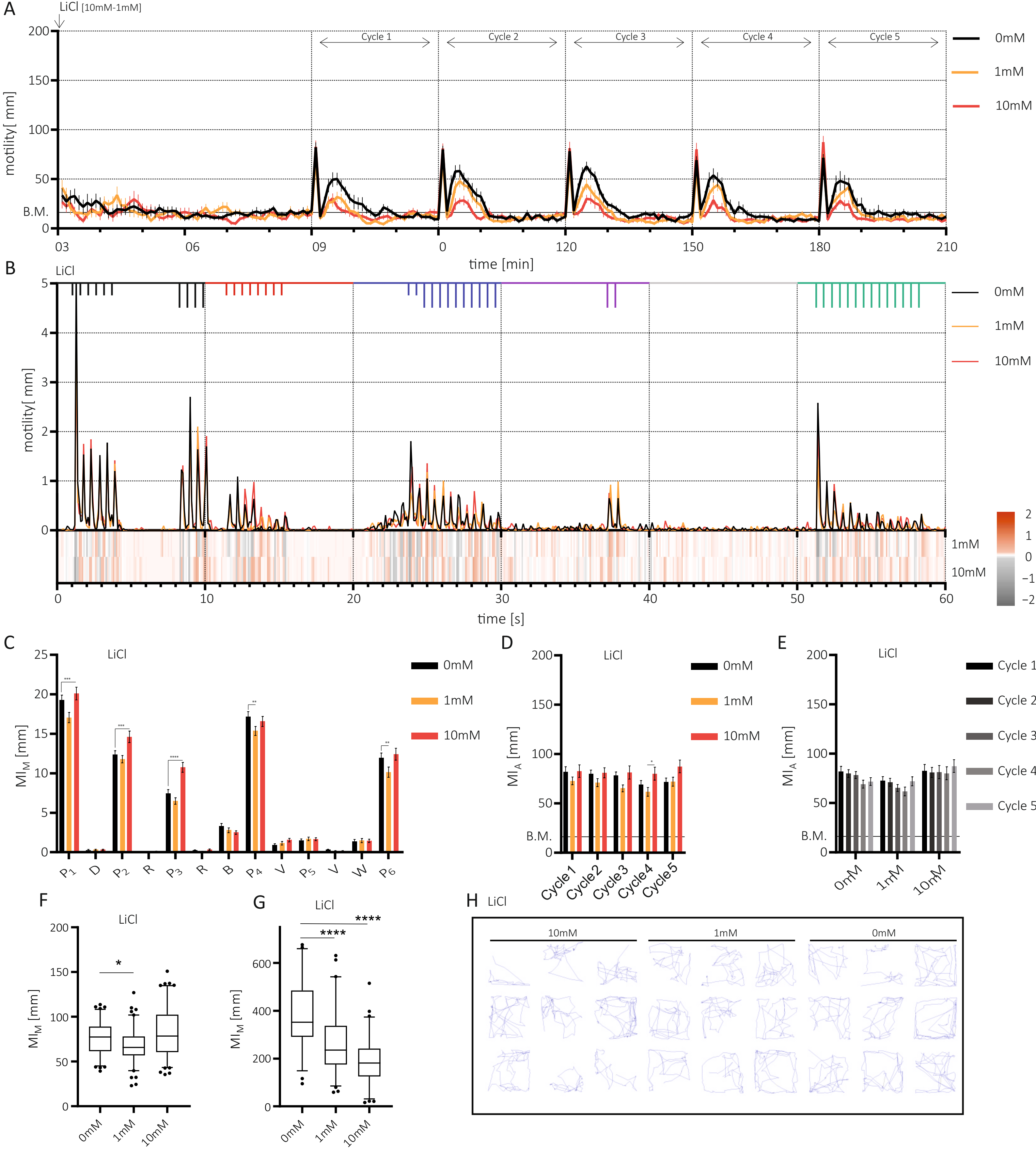


Supplementary figure 4 - Lithium

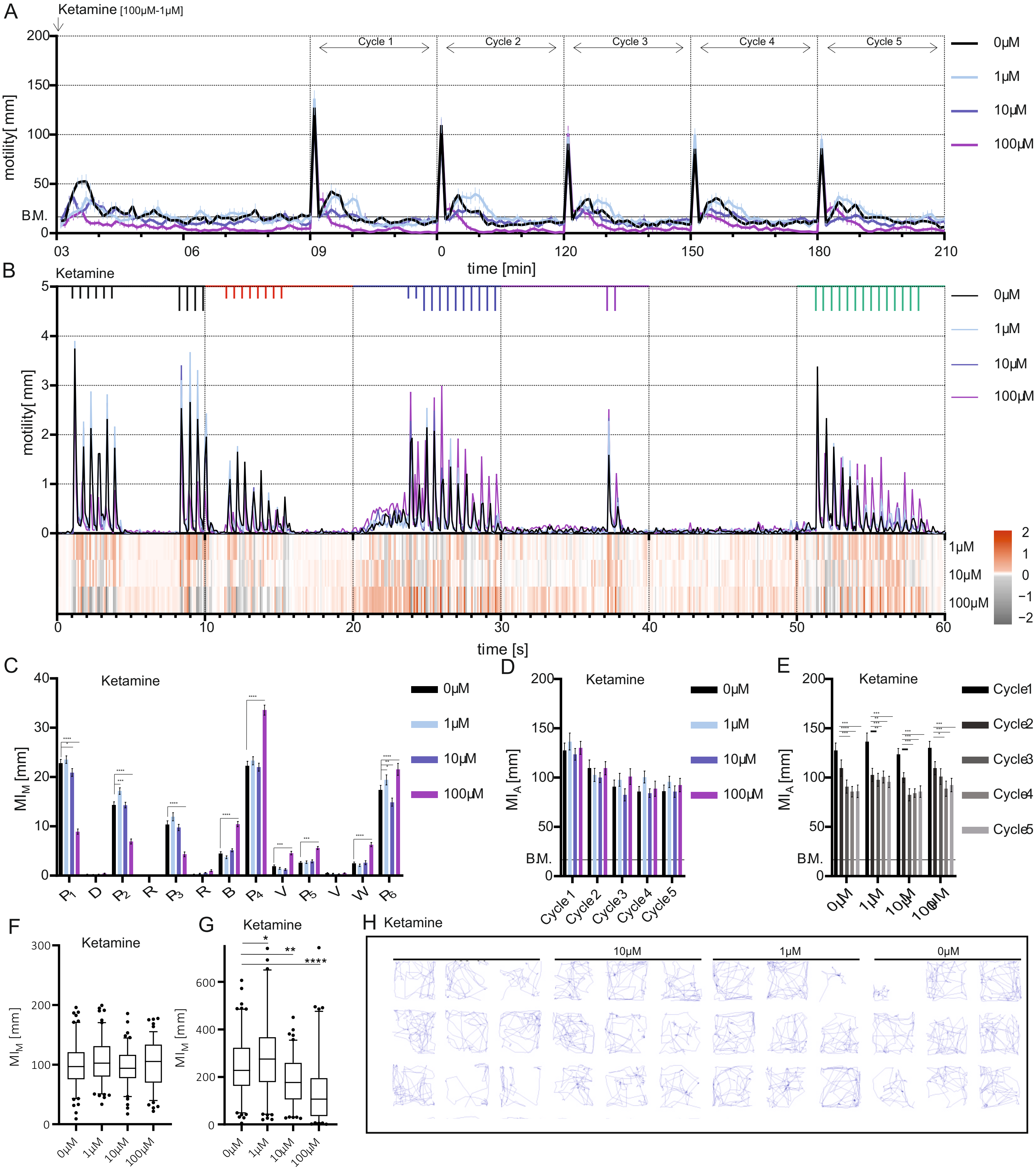


Supplementary figure 5 - Ketamine

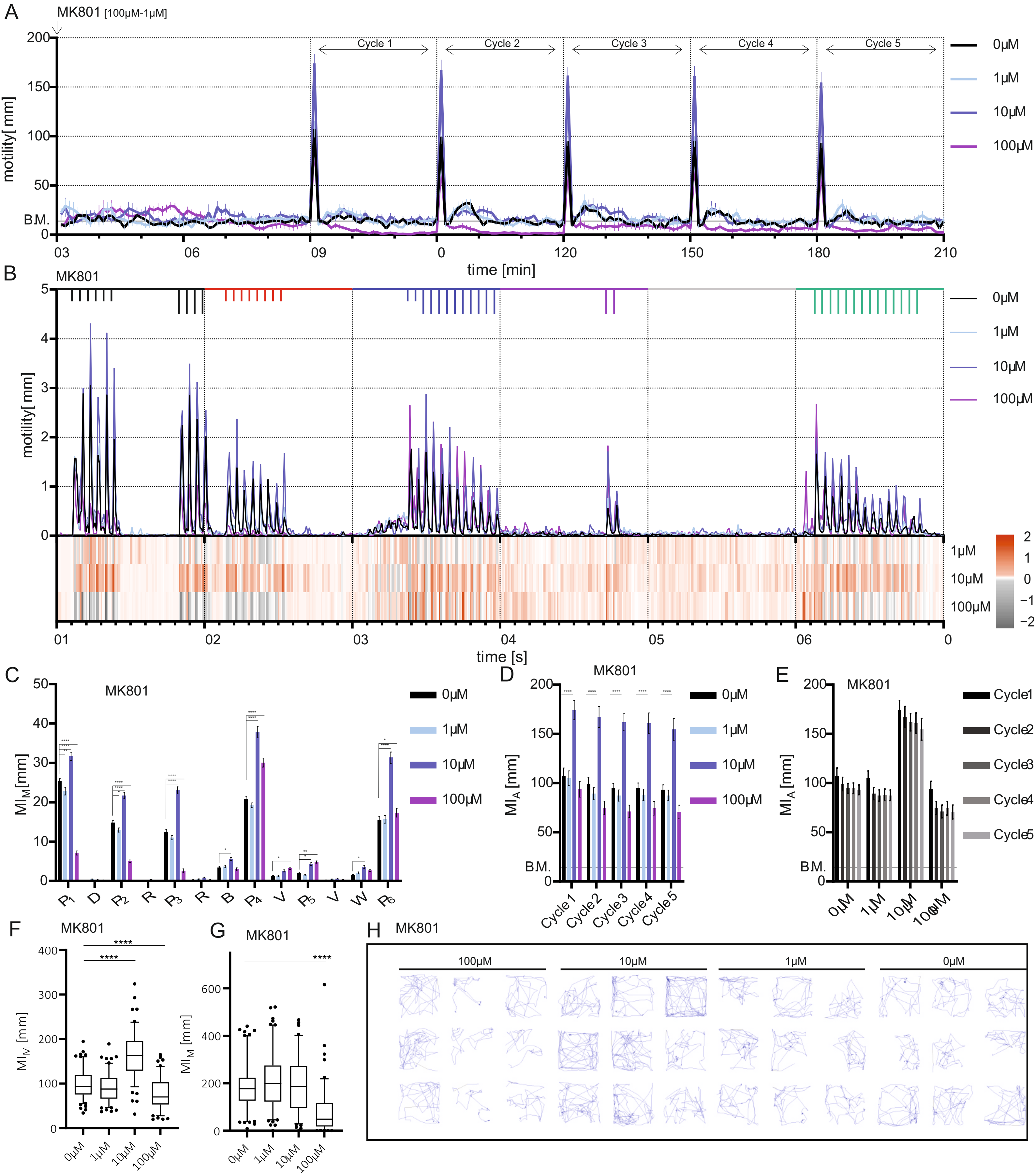


Supplementary figure 6 - MK801

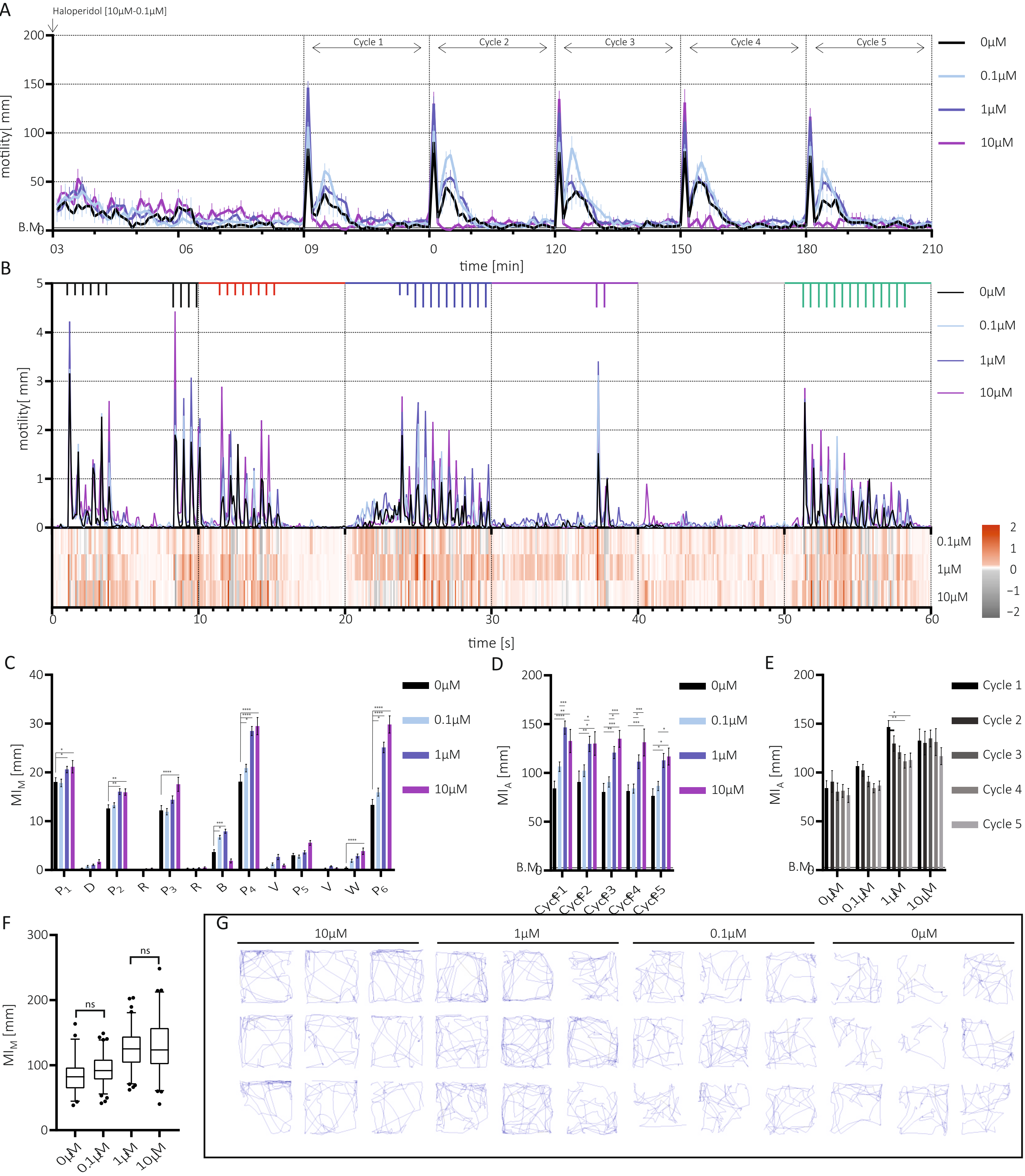


Supplementary figure 7 - Haloperidol

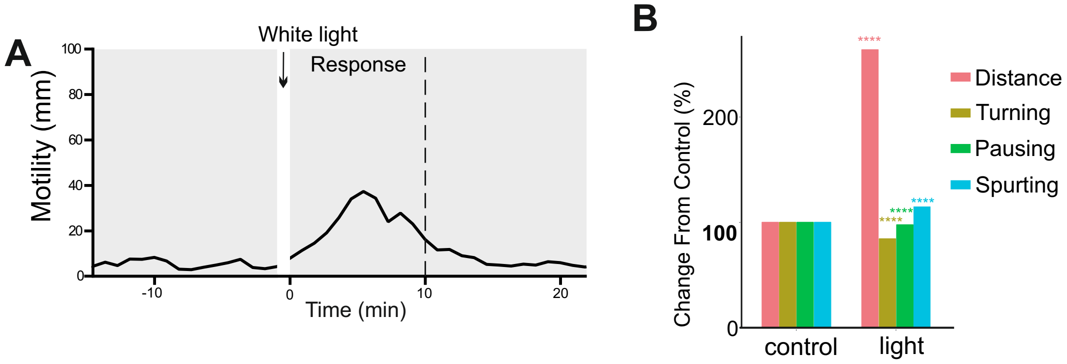


Supplementary figue 8

MATERIALS AND METHODS SUPPLEMENTARY

*Behavioural battery test* - 7 days post fertilization (dpf) zebrafish larvae were transferred to a 96-well plate and drug dosages were administered following which the plate was directly transferred to the DanioVision™ device (Noldus Information Technology, BV, Wageningen, Netherlands) and acclimatized for 1 h in the dark at 28.5 °C. Larvae then underwent the test battery consisting of 5 cycles of a 60 s string of acoustic/vibratory (tapping) and light stimuli programmed using Ethovision13 XT software (Noldus Information Technology, BV, Wageningen, Netherlands). The 60 s stimuli were as follows; high (H) and low (L) intensity patterns (1.6 & 4.2 watt): 1 s pause, 6x L-tap (0.5 s break between individual taps), 4 s pause, 4x H-tap, 1 s pause, 8x L-tap, 8 s pause, 2x L- tap, 10x H- tap, 7 s pause, 2x H-tap, 13 s pause, 14x H-tap, 3 s pause. Visual stimuli evoked using red (635 nm, 192 Lm), blue (470 nm, 240 Lm), purple (red and blue together) and white LEDs, switched colour at 10 s intervals in the following order: darkness, red, blue, purple (red & blue), white, and sequential flickering light of all colours sequentially at 0.1 to 0.2 s intervals. After the stimuli period there was a 29-minute intermission period. This 30 min sequence was repeated 4 more times. Digital video tracking was performed using 60 fps and exported as csv file for analysis using customized software in R.

*Phosphoproteomics* - The brain phosphoproteome data used for this study has been uploaded to the PRIDE database (www.ebi.ac.uk/pride/) to be made open access upon publishing. Reviewers can access this data now using the following credentials: Username:; Password: qNDO1jC9.
